# Organ identity shapes autophagy dynamics and selectivity in plants

**DOI:** 10.64898/2026.07.22.740030

**Authors:** Adrian N Dauphinee, Sanjana Holla, Shamik Mazumdar, Florentine I M Ballhaus, Jonas A Ohlsson, Francis Impens, Teresa Maia, Evy Timmerman, Yasin Dagdas, Christian Löfke, Kerstin Dalman, Karin Schumacher, Peter V Bozhkov, Elena A Minina

## Abstract

Plants, with their unique evolutionary trajectory and complex physiological adaptations, have developed organ-specific mechanisms to cope with various environmental stresses. Autophagy, an essential catabolic process, plays an important role in maintaining plant growth, immunity, and overall fitness. In this study, we reveal the spatio-temporal dynamics of autophagic responses in *Arabidopsis thaliana* roots and shoots under different stress conditions, including AZD8055 treatment and carbon and nitrogen depletion. Our findings demonstrate that roots exhibit stronger autophagic activity than shoots under all three conditions, highlighting the unique adaptations of these two organs. Furthermore, we dissect the selectivity of autophagy in targeting organelles, revealing immediate, delayed, and no uptake categories. Additionally, we observe organelles coexisting within autophagic bodies, shedding light on the complexity of cargo selection. This study enhances our understanding of plant specific autophagy dynamics, emphasizing its role in sustaining the source-sink functions and offering insights into plant adaptation to diverse stressors.

## Introduction

Plant evolution shaped specialized organs responsible for acquiring essential resources: under favorable conditions autotrophic shoots serve as a carbon source for the underground roots, while the roots reciprocate by supplying the necessary nitrogen, potassium, calcium, phosphorus, and other essential minerals. Critically for plant growth and development, these specialized organs must carefully coordinate their functions. For example, nitrogen assimilation is tightly linked to the availability of carbon in the plant, illustrating a delicate equilibrium needed within the plant to optimize resource utilization.^1^ To keep up such balance under conditions where a particular nutrient becomes scarce, plants mobilize the limited resources to prioritize growth of the organ that would best assist in coping with the stress. For instance, under low light conditions that do not aid efficient photosynthesis and result in a scarcity of carbon, plants promote the etiolation of shoots, increasing their chances of reaching a better-illuminated area and re-establish carbon assimilation.^1^ Conversely, when nitrogen availability is low, plants invest into the growth of foraging roots, which search for the nutrient’s sources in the soil.^2^

Such mobilization of inner resources often requires catabolism of existing storages. Autophagy is a powerful catabolic pathway enabling recycling of the cell content to cope with decreased availability of nutrients^3,4^. Its activity is tightly regulated by molecular mechanisms sensing various types of stress and nutrients supply status.

A key regulator of pivotal transition from growth to a stress response in plants is the target of rapamycin kinase complex 1 (TORC1), which monitors nutrient availability. Under optimal conditions, TORC1 fosters plant growth by promoting anabolic processes such as protein synthesis and cell proliferation and suppressing the catabolic activity of autopahgy^5–7^. However, scarcity of nutrients deactivates TORC1, which brings mitosis and translation to a standstill and releases the breaks from autophagy^8^.

The initiation of autophagy sets off a cascade of post-translational modifications that govern the activity of autophagy-related proteins (ATGs) and thereby orchestrate the formation of specialized vesicles in the cytoplasm, known as autophagosomes^9,10^. The autophagosomes envelop various cargoes and traffic them to the lytic compartment, where they are degraded. The resulting breakdown products are then released back into the cytoplasm, supplying the cell with energy and building blocks^10^. Autophagic cargo type range from individual molecules to entire organelles; to aid cell viability, autophagy targets dispensable cell components and must avoid sequestration of the essential ones. Such selectivity of autophagy is thought to be predominantly reliant on the use of autophagy receptors, which label targets available for degradation and even recruit autophagic machinery to construct autophagosomes around them^4,11,12^. Although autophagy observed under conditions of nutrient scarcity is still often referred to as “bulk”, i.e. sequestering cytoplasmic cargo indiscriminately^4,11^, it is worth noting that even in these circumstances, autophagy exhibits a certain degree of selectivity.^13,14^

It is not yet understood how the selectivity of autophagy is suited to keep up the markedly distinct physiology of plant shoots and roots. To address this question, we monitored organ-specific time-resolved changes in activity and selectivity of autophagy when it is deliberately artificially stimulated in both plant organs through a chemical treatment (using a potent selective inhibitor of TORC1 (AZD8055^15^)) and when it was triggered by a physiologically relevant disbalance in the plant macronutrients (**Fig.1**).

**Figure 1.**
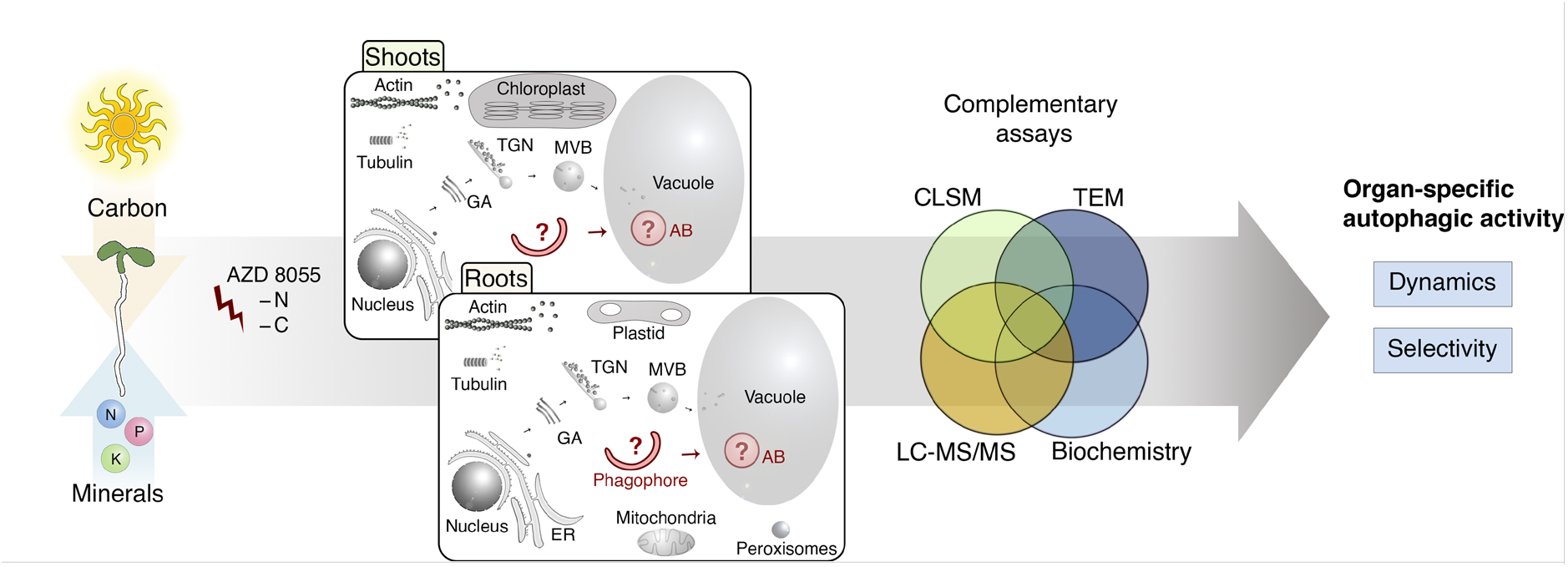
Schematic representation of this study’s objective: organ-specific dynamics and selectivity of plant autophagic activity. Plant organs play distinct physiological roles: while the green photosynthetic tissue serves as a source of carbon, roots are essential for providing the organism with water and minerals. As sessile organisms, plants developed robust stress-coping mechanisms to deal with adverse environmental conditions. One such mechanism is autophagy, a catabolic pathway intricately connected to nutrient sensing and responsive to various stressors. In this study we aim to investigate how autophagy is adapted to the distinct physiology of the plant organs. For this, we plan to investigate organ-specific dynamics of autophagic activity upon treatment with AZD8055, inhibitor of the TORC1, the kinase complex that suppresses autophagy under favorable conditions; and two physiologically relevant stress conditions, i.e. depletion carbon (–C) and nitrogen (–N). We will assess how autophagy impacts cell functionality upon these three conditions by tracking its selectivity towards a representative set of organelles. To ensure robustness of our conclusions, we will implement a combination of complementary methods, including cell biology assays utilizing Confocal Laser Scanning Microscopy (CLSM) and Transmission Electron Microscopy (TEM), biochemical assays, and proteomics (LC-MS/MS).

We aimed to obtain a comprehensive understanding of autophagic activity taking place throughout the entire plant organism: in the autotrophic shoots and heterotrophic roots. For this we deployed a combination of complementary methods, cell biological assays based on confocal and electron microscopy, biochemistry assays and proteomics (**Fig.1**). We revealed clear differences in the timing and range of autophagic response in the roots and shoots and determined dynamic changes in organ-specific selectivity of plant autophagy.

## Results

### Stress-induced plant autophagy exhibits organ-specific dynamics showing a stronger and earlier response in the roots

To consolidate the knowledge already available from the studies on plant autophagy that used either roots or leaves as a model, we first focused on constructing a comprehensive understanding of the timing and extent of the autophagic activity in plant shoots and roots.

For this, we employed AZD8055 (AZD), a selective TORC1 inhibitor which allows to artificially induce plant autophagy with precise timing control^15–17^. The inhibitor was applied to Arabidopsis seedlings expressing autophagy marker pHusion-ATG8A in the wild-type (WT) and autophagy-deficient (*atg5-1/atg7-2*) backgrounds, optimal for monitoring autophagic activity using the previously established Tandem Tag assay^18^ (TT-assay). In brief, the TT-assay relies on autophagy-dependent delivery of pHusion-ATG8A (EGFP-mRFP1-ATG8A) to the lytic vacuole, which results in the accumulation of diffuse red signal in the vacuolar lumen, as the green fluorescence of the tag is quenched by the low pH. Hence, the rising ratio of the red to green fluorescence intensities in cells serves as an indicator for the accelerated pHusion-ATG8 transport to the vacuole, reporting heightened autophagic activity.

Arabidopsis seedlings expressing pHusion-ATG8A were grown, treated with AZD, and imaged within RoPod chambers^19^, yielding time-resolved data on root autophagic activity over a three-day period (**Movie S1**). We observed the formation of numerous autophagosomes in the cytoplasm of roots after 1h of treatment. Their subsequent delivery to the vacuole, resulting in the accumulation of diffuse red fluorescence, was clearly detectable after 2 hours of treatment (**Movie S1**). Therefore, we decided to use 2h of treatment as the earliest time-point of autophagic activity detection in further experiments. Based on the changes in the cell morphology during the treatment, we concluded that viability of root cells was most satisfactory for up to 48h of exposure to AZD. Consequently, we opted to conduct further experiments within this approximate time range.

Next, we proceeded with the time-resolved TT-assay on both plant organs of WT and autophagy-deficient seedlings (**Fig.2A** and **B**, **Fig.S1A**). For each treatment, confocal microscopy was first performed on the roots of complete seedlings mounted inside a RoPod to maintain humidity during imaging, after which cotyledons were detached and used for imaging separately. This approach enabled the near-simultaneous detection of autophagic activity in both roots and shoots within the same biological replicates. As expected, we observed autophagic activity in the root cells after 2h of treatment, however, to our surprise, the response in shoots was detectable only after 24h of treatment (**Fig.2A** and **B)**. Furthermore, the strongest observed autophagic activity was nearly two-fold higher in roots compared to shoots (**Fig. 2B**). Interestingly, we also detected a decrease in autophagic activity after 56h of treatment, followed by slight increase at 72h. These results indicate a possibility of an oscillating plant autophagic response during prolonged stress, similar to what has been previously reporter for animal models^20,21^ (**Fig2.B**).

**Figure 2.**
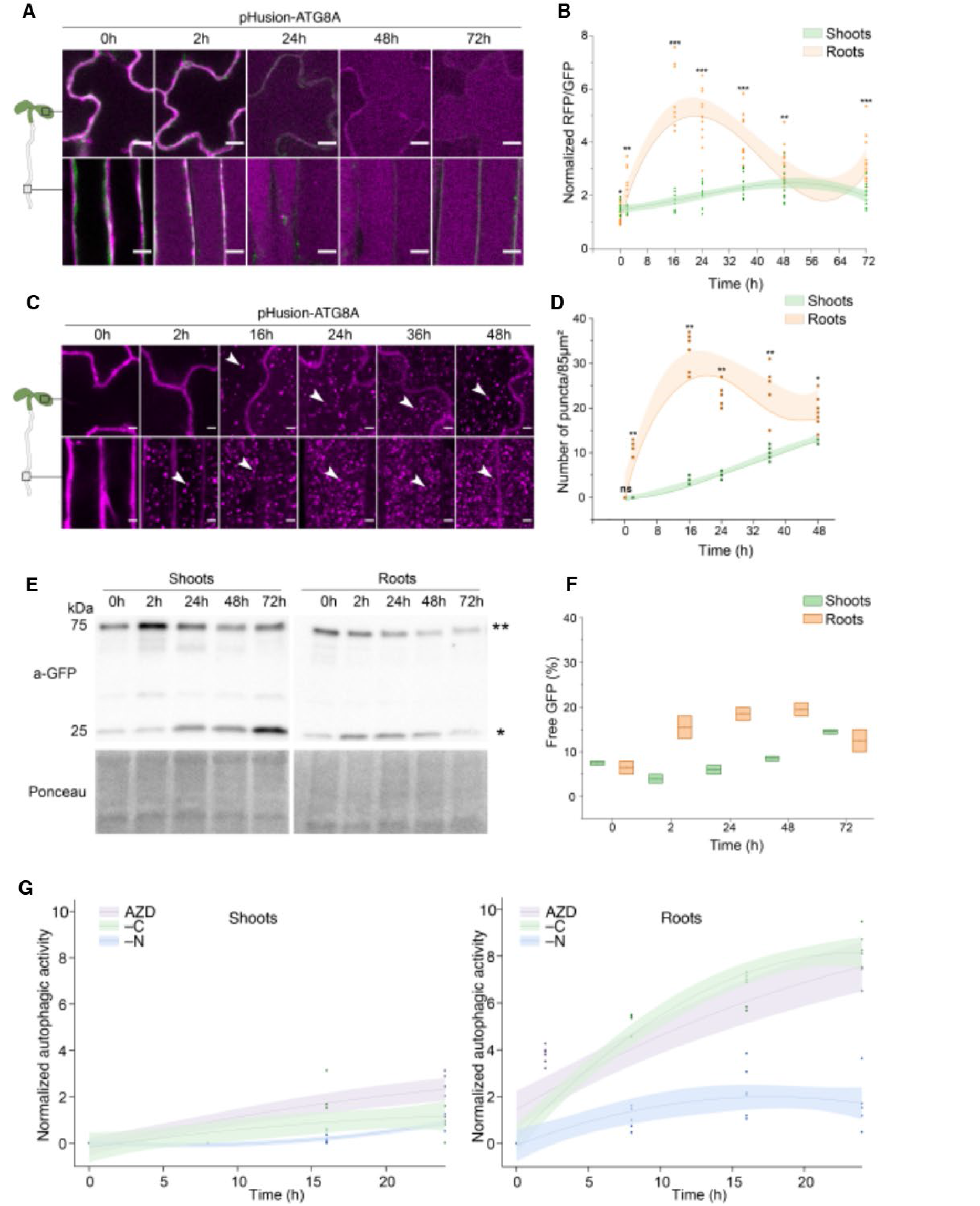
All three stress conditions elicit stronger and earlier autophagic response in roots. **A.** Tandem-tag assay reveals the organ-specific autophagic response to AZD8055 (AZD): 5-days-old *Arabidopsis thaliana* seedlings expressing pHusion-ATG8A (EGFP-mRFP1-ATG8A) were treated with AZD for the specified durations prior to imaging using CLSM. Fluorescent marker is delivered to the root vacuoles already upon 2 hours of treatment and only upon 24 hours of treatment to shoot vacuoles. Localization of the same marker in autophagy-deficient *atg5-1/atg7-2* seedlings is shown in the (**Fig.S1**). Scale bars, 10 μm. **B.** Quantitative analysis of organ-specific autophagic response illustrated in (**A**). Autophagic activity over time was detected by quantifying the ratio of red to green fluorescent signals within the vacuoles. The ratios for autophagy-deficient *atg5-1/atg7-2* seedlings were used as a background to normalize the corresponding values obtained on the WT seedlings. Autophagic activity increases in roots already after 2 hours of treatment, whereas the shoots respond only after 24 hours. The highest detected activity in the two organs differs almost two-fold. Data represents three independent experiments. n=91, Unpaired two-tailed Student’s t-test, *, p-value<0.05; **, p-value<0.01; ***, p-value<0.001. Trendlines represent polynomial fit; shaded regions denote 95% confidence bands. **C.** Confocal microscopy performed on 5-days-old pHusion-ATG8A seedlings treated with AZD and Concanamycin A (ConA) shows accumulation of autophagic bodies in the root vacuoles already after 2 hours of treatment and only after 24 hours in the shoot vacuoles. Only the red channel shown. White arrowheads indicated autophagic bodies inside the vacuoles. Scale bar, 10 μm. **D.** Quantification of vacuolar puncta illustrated in (**E**) shows higher accumulation of autophagic bodies in the roots when compared to shoots, thereby corroborating the organ-specific autophagic response observed in (**A**-**D**). Unpaired two-tailed Student’s t-test, *, p-value<0.01; **, p-value<0.001; ns- not significant. Data represents two independent experiments, n=108. **E.** GFP-cleavage assay was performed on shoots and roots harvested from pHusion-ATG8A seedlings treated with AZD for the indicated amount of time. **, Full-length pHusion-ATG8A; *, cleaved EGFP. Ponceau staining was used as loading control. **F.** Quantification of cleaved EGFP from the pHusion-ATG8A corroborates organ-specific autophagic response shown in (**A**) and (**B**). Data represents two independent experiments. Median is indicated by a line across the box. Mean is indicated by the circle on the median. **G.** Representative data of autophagic activity in shoots and roots of 5-days-old seedlings expressing mCherry-ATG8A that were treated with AZD, nitrogen (–N) or carbon (–C) depletion, in the presence of ConA. Accumulation of autophagic shows that autophagic activity is higher in roots in comparison to the shoots under all three experimental conditions. Trendlines represent polynomial fit; shaded regions denote 95% confidence bands.

A prolonged exposure to AZD inhibits protein synthesis and enhances protein degradation, which eventually depletes the fluorescent autophagy reporter^19^ and might lead to suboptimal quality of the data. Therefore, we decided to corroborate the results obtained with the TT-assay by also performing puncta quantification assay in the seedlings co-treated with AZD and Concanamycin A(ConA), where the latter prevents degradation of the autophagosomes delivered to the vacuoles^22^. This method revealed presence of autophagic activity in the shoots at an earlier time point, already upon 16h of treatment (**Fig.2C** and **D**). This might be attributed to the fact that weak fluorescence signals are more efficiently detected when they are concentrated in the form of puncta, rather than being diffusely present in the large volume of the vacuolar lumen. Nonetheless, it was consistently evident that the autophagic response in the shoots was significantly delayed in comparison to the roots (16h *vs* 2h) (**Fig.2C**). Additionally, this assay confirmed that autophagic activity in roots is nearly two-fold higher compared to the maximal activity observed in the shoots (**Fig.2C**).

The above-described experiments rely on confocal microscopy restricted to epidermal cells of the corresponding organ. To validate that our findings can be extrapolated on the entire plant organ, we performed GFP-cleavage assay^22^ on dissected shoots and roots of seedlings treated with AZD (**Fig.2E** and **F**). A small amount of free GFP was observed in each organ under non-inducing condition, representative of the basal autophagic flux. In agreement with our results, accumulation of the free GFP was significantly delayed in the shoots compared to roots and the efficiency of GFP cleavage from the pHusion-ATG8A was higher in the roots (**Fig.2E** and **F**). No processing of the fusion protein was observed in the autophagy-deficient seedlings subjected to the same treatment, corroborating that the process is autophagy-dependent (**Fig.S1B**).

Primary roots with developed root hairs are plausibly more efficient in absorbing compounds from the medium than the shoots. Therefore, we questioned whether the organ-specific timing of the autophagic response to AZD treatment was reflective of the inherent differences in autophagy regulation in these organs or occurred due to disparities in the compound uptake efficiency. First, we assessed cell permeability for three small organic molecules commonly used in plant biology and with molecular weights similar to AZD (MW=465Da): fluorescein diacetate (FDA, MW=416 Da); propidium iodide (PI, MW=668 Da); and the styryl dye FM4-64 (MW=664 Da). Upon a short incubation, each of these three fluorescent compounds could be detected in both organs within only a few minutes of delay for the shoots (**Fig.S2**). These results indicated that 14-22h delay in the autophagic response of the shoots is not caused by slower uptake of AZD in this organ.

Furthermore, we hypothesized that AZD uptake should be most impacted by the organ architecture, cuticle layer and cell walls. We removed these potential barriers by isolating protoplasts from roots and shoots and then subjecting them to AZD/ConA treatment. Isolated cells of shoots and roots exhibited the same response timing as we previously observed in the corresponding intact organs (**Fig.S3A**). These results demonstrated that the differences in the timing of autophagic response of shoots and roots were intrinsic to the cells of these organs.

Since cotyledons are a specialized type of leaves^23^, we considered it important to verify that our observations obtained on the seedlings’ leaves are representative for the plant shoot. For this, we exposed the true leaves of 5-week-old Arabidopsis plant to AZD/ConA treatment for 0h, 2h and 24h and indeed observed a detectable response only at 24 of treatment (**Fig.S3B**). Additionally, mesophyll protoplasts from the leaves of the same plants when exposed to the same treatment also exhibited autophagic activity only after 24h (**Fig.S3C**). These results confirmed that indeed AZD-induced autophagy is detectable in the plant shoots upon approximately one day of treatment. We were not able to develop a reasonable approach to reliably monitor AZD-triggered autophagic response in the root system of soil-grown plants and set aside this question for future research.

Arabidopsis genome contains nine orthologs of ATG8 (ATG8A-I isoforms), which might be playing at least partially distinct functions in Arabidopsis autophagy^24,25^. All of the above-described experiments were carried out using the ATG8A isoform. To verify that these observations are representative of Arabidopsis autophagy we detected autophagic response in shoots and roots using each of the nine ATG8 isoforms. Notably, all ATG8 isoforms reported a very similar response to the AZD, showing earlier and stronger autophagic activity in the roots (**Fig.S4**).

Finally, we endeavored to test the organ-specific plant autophagic response to the stress triggers that are more physiologically relevant. For this we subjected seedlings to depletion of macronutrients: carbon (–C) or nitrogen (–N). We considered comparison of organ-specific response to these two commonly used inducers of plant autophagy as most interesting, because carbon depletion promotes shoot etiolation and stagnates root growth ^26^, whereas depletion of nitrogen results in the opposite phenotype by slowing down the shoot growth and promoting root growth^27^. Surprisingly, despite the opposite effects on the plant organ growth, both –C and –N elicited earlier and stronger autophagic response in the roots than in shoots (**Fig.2G**, **Fig. S5**).

In summary, we performed a thorough validation of time-resolved autophagic activity occurring in roots and shoots of Arabidopsis seedlings subjected to AZD, –C and –N treatments. Our findings consistently demonstrated a stronger and earlier autophagic response that is inherent to root cells.

### AZD induces simultaneous occurrence of selective and bulk autophagy in both plant organs

The distinct timing and extent of autophagic response in shoots and roots led us to question what its intended targets might be. Firstly, we assessed whether autophagy in either of the organs exhibits selectivity or takes up cargo indiscriminately. We hypothesized that it would be highly unlikely for plants to possess autophagy receptor recognizing mRFP, a fluorescent protein from an aquatic animal^28^, and therefore it can be deployed as a reporter for receptor-independent sequestration of soluble cytoplasmic proteins into Arabidopsis autophagosomes, i.e. non-selective autophagy. We exposed seedlings expressing pHusion-ATG8A or mRFP to AZD/ConA treatment for 0-48h and monitored accumulation of autophagic bodies in the vacuoles of shoots and roots (**Fig.3A**). mRFP was detectable in the vacuolar puncta of roots and shoots immediately upon initiation of autophagy in the corresponding organ, i.e. at 2h in roots and at 16h in shoots **(Fig.3A)**. These results suggested that in both plant organs AZD induces autophagy which unselectively takes up soluble cytoplasmic proteins, the so-called “bulk autophagy”^4,12^.

**Figure 3.**
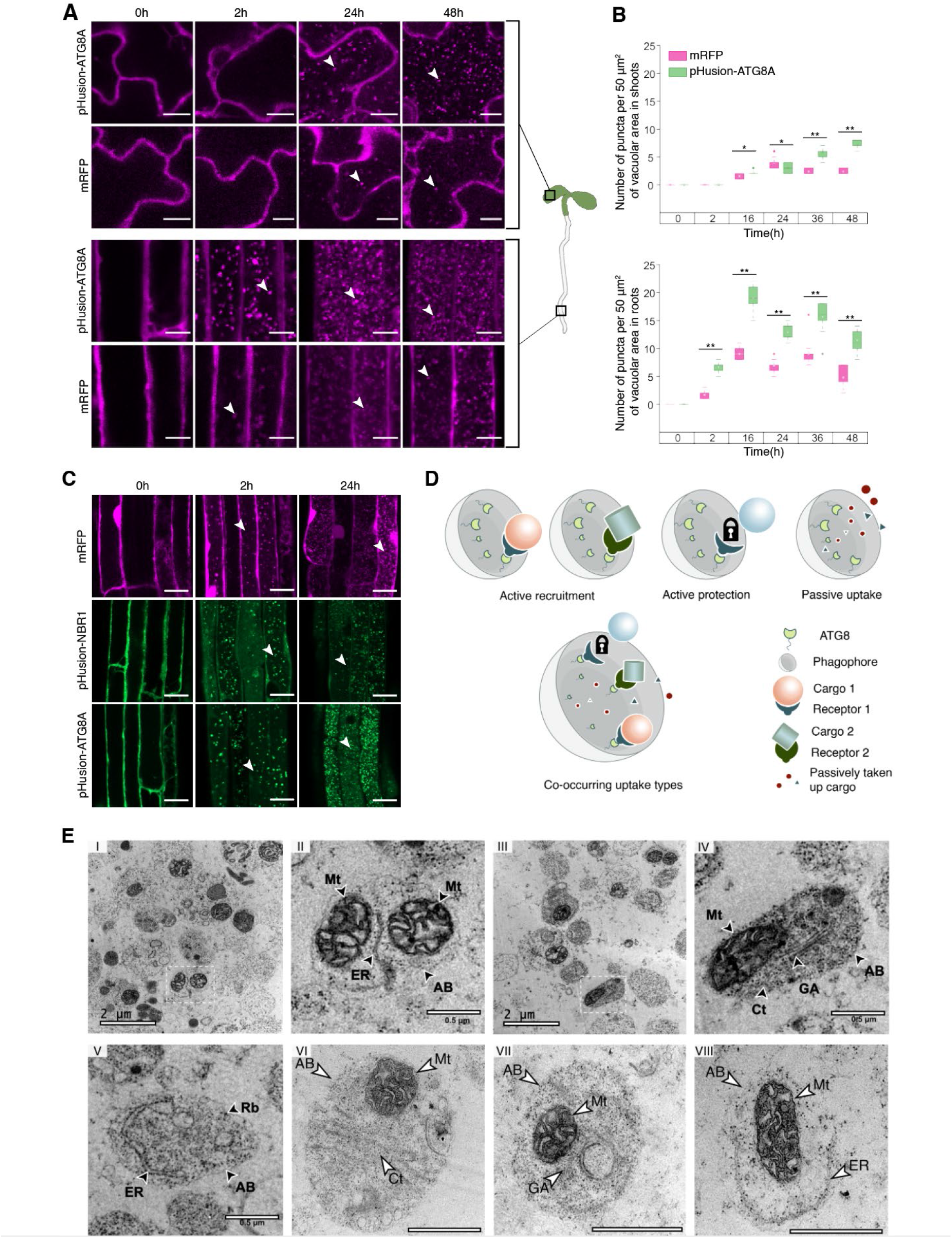
AZD simultaneously triggers receptor-dependent and independent autophagy. **A.** Confocal microscopy performed on 5-days-old Arabidopsis seedlings expressing pHusion-ATG8A or mRFP (reporter for passive cargo uptake by autophagosomes) treated with AZD/ConA for the specified time durations. Similar to pHusion-ATG8, mRFP is detectable in autophagic bodies after 2h of treatment in the roots and after 24h in the shoots. White arrowheads indicate autophagic bodies in the vacuoles. Scale bars, 10 µm. **B.** Quantitative analysis of the data illustrated in **A** reveals lower autophagic activity in the shoots compared to roots detectable with either of the markers. Data represents three independent experiments, n=90, unpaired two-tailed Student’s t-test, *, p-value<0.05; **, p-value<0.001; ***, p-value<0.0001. Median is indicated by a line across the box; mean is indicated by small circles; range within 1.5IQR is represented by whiskers; outliers are represented by solid rhombus. **C.** Confocal microscopy performed on the roots of 5-days-old Arabidopsis seedlings expressing pHusion-ATG8A, pHusion-NBR1 (reporter for receptor-dependent cargo uptake by autophagosomes) and mRFP. Seedlings were with AZD/ConA for the indicated time durations. All three markers are delivered to the vacuole after 2 hours of drug treatment. White arrowheads indicate autophagic bodies in the vacuoles. Scale bars, 20 μm. **D.** Schematic depiction of potential mechanisms for sequestering cargo into autophagosomes: (i) receptor-dependent active recruitment. In this process, specific receptors actively target corresponding cargo for sequestration into autophagosomes, possibly involving interaction with ATG8; (ii) active protection, wherein either cargo is masked from its receptor, or the receptor-cargo complex is not recognized by ATG8; (iii) passive cargo uptake, which might be represented by diffusion of soluble cytoplasmic proteins into the forming autophagosome; (iv) co-occurring cargo uptake, formation of autophagosomes may employ a combination of the above-described cargo recruitment strategies. **E**. TEM micrographs obtained on Arabidopsis roots expressing pHusion -ATG8A in the WT background and treated with AZD/ConA for 24 hours reveal that plant autophagic bodies contain a mix of various cargo types. i. Multiple autophagic bodies observed in the vacuole of a root epidermal cell. ii. Zoomed-in selection indicated in (i) shows a single autophagic body containing two mitochondria and a portion of ER. iii. Overview of autophagic bodies accumulated in another epidermal cell of the same root. iv. Zoomed-in selection indicated in (iii) shows a single autophagic body containing a mitochondrion, GA stack and a portion of cytoplasm. v. Autophagic body containing ER strand and portion and cytoplasm with ribosomes. vi. A micrograph of an autophagic body containing a mitochondrion, cytoplasm and potentially ER. vii. A micrograph of an autophagic body containing a mitochondrion, a GA stack and cytoplasm. viii. A micrograph of an autophagic body containing a mitochondrion, portion of ER and cytoplasm. AB, autophagic body; ER, endoplasmic reticulum; Mt, mitochondria; GA, Golgi apparatus; Rb, ribosomes; Ct, cytoplasm. For vi-viii scale bars, 1μm.

However, quantification of the autophagic bodies in the vacuoles of shoots and roots revealed a significantly higher number of pHusion-ATG8A-labelled puncta compared to mRFP-positive puncta **(Fig.3B)**, indicating a simultaneous formation of strictly selective autophagosomes that exclude the receptor-independent cargo and “bulk” autophagosomes that include it. Interestingly, the number of mRFP-positive puncta in the vacuoles of shoot cells reached a plateau at 24h of treatment, while the number of pHusion-ATG8A-positive puncta continued to rise (**Fig.3B**). This suggests an increase in autophagy selectivity upon prolonged stress in the shoot. Notably, such an increase in autophagy selectivity was not observed in the roots (**Fig.3B**).

We corroborated occurrence of selective autophagy under AZD by monitoring inclusion of NBR1 into autophagic bodies upon AZD/ConA treatment. NBR1 is a well-characterized autophagy receptor for ubiquitinated cargo^29^. Indeed, we could observe simultaneous formation of NBR1-positive and mRFP-positive autophagic bodies immediately upon activation of autophagy (**Fig.3C**).

Based on the current knowledge on cargo sequestration into autophagosomes^4,12^ and our observations, we proposed that plant autophagy can exhibit such complex selectivity using four possible scenarios (**Fig.3D**): (i) active recruitment of the cargo, executed by receptors recognizing specific targets and associating with the growing phagophore *via* ATG8; (ii) active protection of essential cellular components by masking them from the corresponding autophagy receptors or masking the receptor-cargo complex from interaction with ATG8; (iii) passive uptake of a cargo that diffuses into the forming autophagosome and gets trapped in the vesicle upon its completion; (iv) co-occurring cargo uptake, during which all of the three above-listed strategies happen simultaneously.

A recent study demonstrated that in starved *Saccharomyces cerevisiae* cells most autophagosomes sequester ribosomes-containing cytoplasm by a mechanism most resembling “passive uptake”, while a few autophagosomes additionally contain specific types of cargo, for example amorphous Ede1-dependent endocytic protein deposit surrounded by ER^13^, indicating that our suggested “co-occurring uptake” strategy is indeed executed by yeast autophagy. To test whether plant autophagosomes can be formed in a similar manner, we performed TEM on the WT and autophagy-deficient Arabidopsis seedlings treated with AZD/ConA for 24h. Imaging of epidermal root cells revealed accumulation of numerous electron-dense vesicular structures in the WT vacuoles which were not present in the vacuoles of the *ATG*-knockouts (**Fig.S6**), therefore we concluded that these structures are autophagic bodies. A closer analysis of the autophagic bodies content demonstrated that they contain a mix of various cargo types (**Fig.3E**): a combination of mitochondria, ER and cytoplasm (**ii** and **viii**); or mitochondria, Golgi apparatus stack and cytoplasm (**iv** and **vii**). Consistent with the presence of the non-selective marker (mRFP) in vacuolar puncta (**Fig.3A** and **B**), we could see cytoplasm in most if not all detected autophagic bodies (**Fig.3E**). In summary, the ultrastructural imaging corroborated occurrence of receptor-independent “passive cargo uptake” and revealed presence of several types of cargoes in the same autophagosome.

Since a TEM section represents only a very thin portion of each autophagic body it was not possible to decidedly conclude whether we could observe any autophagic bodies containing exclusively one type of cargo. Furthermore, not all types of organelles could be easily discerned on TEM images without additional labelling. Therefore, to further explore potential selectivity during AZD-induced plant autophagy we deployed CLSM using fluorescent markers for a representative set of plant organelles.

### Plant autophagy exhibits organ-specific prioritizing in targeting cell components

To investigate what cellular components are targeted by autophagy in the two plant organs, we established a set of WT and autophagy-deficient Arabidopsis lines co-expressing autophagy reporter mCherry-ATG8A with GFP-tagged markers for membrane organelles.

We treated Arabidopsis seedlings with AZD/ConA for 48h and monitored autophagy-dependent delivery to the vacuole of six organelles (endoplasmic reticulum, ER; Golgi apparatus, GA; trans-Golgi network, TGN; multivesicular bodies, MVBs; peroxisomes, and mitochondria), two plant cytoskeleton components (tubulin and actin), and cytoplasm (mRFP) in shoots and roots (**Fig.4**, **Fig.S7**). Remarkably, while some cell components were immediately sequestered into autophagosomes upon activation of autophagy in both plant organs, targeting others was clearly differently prioritized in the shoots and roots (**Fig.4**, **Fig.S7**). For instance, ER was present in the autophagic bodies of roots and shoot as soon as autophagic activity became detectable in the corresponding organs. Conversely, GA was detectable within autophagic bodies immediately upon upregulation of autophagy in the shoots, but it was taken up by autophagosomes in the roots only after a prolonged exposure to AZD (22h after autophagic activity became detectable) (**Fig.4A** and **B**). This indicates that in roots GA might be temporarily selectively protected from being degraded, and this protection is lifted if cells are exposed to a prolonged stress. Furthermore, peroxisomes were not detectable within autophagic bodies throughout the whole duration of treatment (**Fig.4A** and **B**), illustrating that some cell components are better protected than others. Moreover, we observed that certain organelles, such as the ER and MVBs, were present in approximately 50-75% of the autophagic bodies **(Fig. 4B; Fig. S7B)**, meaning that they must co-occur withing the some of the autophagic bodies. This provides additional evidence to our observations obtained with TEM that under AZD treatment, plant autophagosomes sequester a mix of different cargo types.

**Figure 4.**
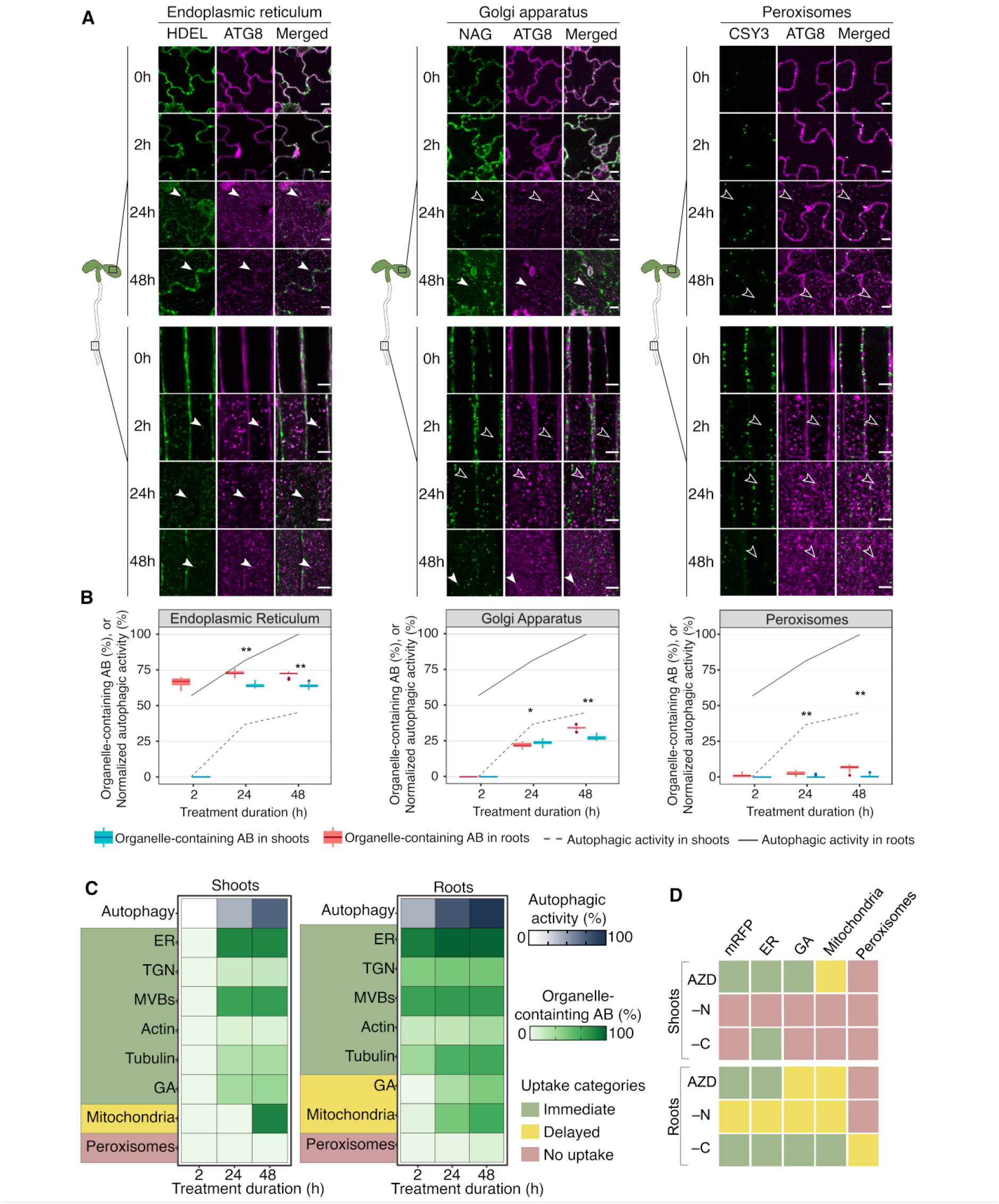
Plant autophagy in shoots and roots differs in its prioritization for targeting cell components. Eight markers for organelles and cytoskeleton were checked for their sequestration into autophagosomes in shoots and roots of Arabidopsis seedlings. Based on the timing of their uptake into the autophagosomes, the organelles were categorized into three groups: immediate uptake, delayed uptake and no uptake. (**A**) and (**B**) display data for one representative organelle from each of these groups. **A.** Confocal microscopy images obtained on 5-days-old Arabidopsis seedlings co-expressing mCherry-ATG8A with one of the following organelle markers: GFP-HDEL (endoplasmic reticulum), NAG-GFP (Golgi apparatus) and CSY3-GFP (peroxisomes). The seedlings were treated with AZD/ConA for the indicated amount of time. For roots, the endoplasmic reticulum, Golgi apparatus, and peroxisomes were placed into immediate uptake, delayed uptake, and no uptake categories, respectively. In the shoots, the Golgi apparatus was detected as the immediate uptake category. Filled arrowheads indicate autophagic bodies containing an organellar marker, outlined arrowheads point at autophagic bodies without the corresponding organellar marker. Scale bars,10 µm. **B.** The charts represent autophagic activity detected in the shoots (dotted line) and in the roots (solid lines), along with the percentage of autophagic bodies containing the corresponding organelle (box plots). Data represents two independent experiments. n =72, unpaired two-tailed Student’s t-test, *, p-value<0.01; **, p-value<0.0001. **C.** The heat map represents autophagic activity (blue) and sequestration rate of organelles detected in roots and shoots, at three time points of AZD/ConA treatment. The heatmap is built based on the data presented in Fig.4B and **Fig. S7B**. **D.** Heatmap illustrating the three categories of organelle uptake observed in the shoots and roots under AZD treatment, nitrogen (–N) and carbon (–C) deprived conditions. The mRFP was used as a marker for passive cargo sequestration. ER, Endoplasmic reticulum; GA, Golgi apparatus; TGN, trans-Golgi network; MVBs, multivesicular bodies. The heatmap is built based on the data shown in Fig. 4B, **Fig. S7B**, **Fig. S8B** and **Fig. S9B.** Data represents two independent experiments for each condition.

We determined that ER, GA, TGN, MVBs, ACTIN and TUBULIN were taken up into the autophagic bodies of shoots at the earliest time point with detectable autophagy (24h), mitochondria were targeted only later (at 48h); whereas peroxisomes remained protected throughout the treatment (**Fig. 4A-C**; **Fig.S7A** and **B**). In roots, ER, TGN, MVBs, ACTIN and TUBULIN were immediately targeted by autophagy (detectable in the autophagic bodies after 2h of treatment), GA and mitochondria were sequestered into autophagosomes only on the 24th hour of stress, while peroxisomes remained protected from degradation (**Fig. 4A-C**; **Fig.S7A** and **B**). The markers of cellular components were not detectable in the vacuolar puncta when expressed in the autophagy-deficient backgrounds (**Fig.S7C**), confirming that their uptake is autophagy dependent.

Based on the timing of cell component marker sequestration into autophagosomes, we derived three categories for their uptake by autophagy: “immediate uptake” for cell components targeted as soon as autophagy is initiated; “delayed uptake” for those sequestered into autophagosomes significantly later, and “no uptake” for components protected from degradation (**Fig. 4C**). According to this classification, we determined that GA is more protected in the roots than in shoots, despite roots exhibiting a much stronger autophagic activity.

In summary, these results confirm our above-described observation of the co-occurring selective and receptor-independent autophagy under AZD treatment, resulting in sequestration of several cargo types by at least some autophagosomes. Furthermore, they demonstrate that selectivity of autophagy is organ-specific and changes under prolonged stress allowing targeting a broader range of cargo types.

To gain further insight into the organ-specific targeting of cell components, we subjected reporter lines expressing markers for ER, GA, mitochondria, peroxisomes, and cytoplasmic soluble proteins (mRFP) to –N and –C (**Fig.4D**, **Fig.S8**, **Fig.S9**). Due to cytotoxic effect of –N/ConA and –C/ConA treatments, we limited the duration of the experiments to only 24h, during which cells of shoots and roots exhibited standard morphology.

–C treatment caused immediate uptake of ER, GA, mitochondria, and mRFP into the autophagosomes of root cells, and delayed uptake of peroxisomes. Remarkably, in the shoots of the same seedlings, only ER was targeted by autophagy, while the other markers, including cytoplasmic mRFP, were protected from degradation **(Fig.4D, Fig. S8)**. These results demonstrate a stricter selectivity of autophagy in the shoot compared to root under carbon depleted conditions.

Most intriguingly, the depletion of nitrogen yielded even more selective autophagic response: although autophagy was detectable in the roots already at 8h of AZD/ConA treatment, none of the markers for cell components, were observed in the vacuolar puncta, not even the cytoplasmic mRFP (**Fig.S9**). However, eight more hours of treatment eventually triggered uptake of mRFP and ER by autophagy, while GA and mitochondria were targeted with an even bigger delay, at 24h of treatment (**Fig.4D, Fig.S9**). The earlier sequestration of the passive uptake marker than of GA and mitochondria (**Fig.S9**) suggests that the latter might indeed be actively protected from recruitment into autophagosomes (**Fig.3D**). Notably, root cell peroxisomes remained protected from autophagy throughout the duration of –N treatment (**Fig.4D, Fig.S9**). In the shoots, despite the presence of autophagic bodies at 24 hours, none of the cell components markers could be detected in the vacuolar puncta (**Fig.4D, Fig.S9**).

In summary, under both –C and –N conditions we observed higher selectivity of autophagy in the shoots compared to roots. Furthermore, the absence of the cytoplasmic marker (mRFP) in autophagic bodies under these conditions suggests that elongation of the forming autophagosomal membrane was carried out in the close proximity to the cargo, thereby preventing passive diffusion of soluble cytoplasmic components into the forming vesicle.

### Vacuolar proteomes reveal organ-specific cargo targeting by autophagy

We sought to expand our observations beyond the use of fluorescent markers, which allow to monitor autophagic targeting of only individual organelles, and predominantly in the epidermal cells, and aimed to assess representativeness of our results for the entire corresponding plant organ. For this we focused on obtaining quantitative vacuolar proteomes that would allow us to identify in a high-throughput manner proteins that are delivered to the vacuoles of shoots and roots upon upregulation of autophagy.

To achieve this, we developed a protocol that would permit extracting total proteins from isolated plant cells and from intact vacuoles upon 2h and 24h of autophagy induction with AZD/ConA, with the aim to later use them for isobaric labelling followed by quantitative LC-MS/MS (**Fig.5A**). Resulting proteomes can be then compared to identify proteins that became enriched in the vacuoles of shoots or roots upon induction of autophagy and determine what proportion of each protein was delivered to the vacuole by autophagy (**Fig.5A**). First, we established the protocol for intact plant vacuole extraction under normal and autophagy-inducing conditions. Using seedlings expressing green fluorescent cytoplasmic marker and red fluorescent marker for vacuolar lumen, we could confirm isolation of intact vacuoles (**Fig.S10A**) with untraceable amounts of cytoplasmic contamination (**Fig.S10B**). Next, we verified that autophagy could be reproducibly induced in the large amounts of shoot and root protoplasts required for subsequent extraction of the vacuoles, and that it yields the response comparable with our previous observations, i.e. autophagic response was detectable in the root cells after 2h and in the shoot cells after 24h of induction (**Fig.S10C**). Finally, we ensured that we can harvest sufficient amount of protein required for the subsequent mass spectroscopy by performing total protein extraction from complete protoplasts subjected to autophagy induction for 0h, 2h and 24h and from their corresponding vacuolar fractions (**Fig. S10C**). Expectedly, total protein content of shoot protoplasts was significantly higher compared to the roots, and vacuolar fractions contained less protein than whole-cell extracts (**Fig.S10C**). Furthermore, prolonged AZD/ConA treatment was causing a significant decrease in the protein content in the samples (**Fig. 10C**).

**Figure 5.**
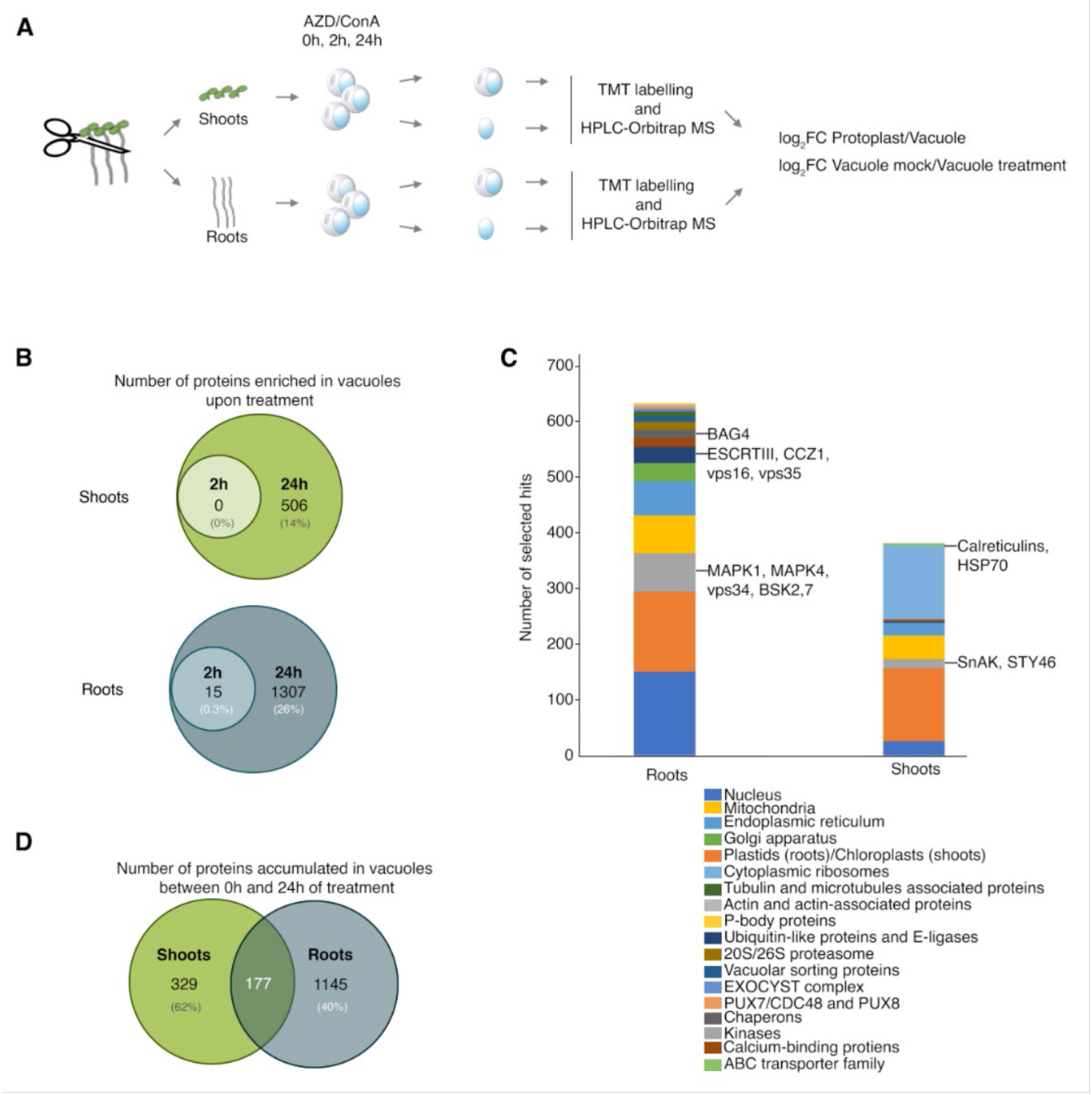
Analysis of proteome changes in root and shoot vacuoles upon autophagy induction. **A.** Schematic representation of the quantitative proteomics analysis. Seven days-old seedlings grown under standard conditions were used to isolate protoplasts from shoots and roots. Isolated cells were then subjected to AZD/ConA treatment for 0h, 2h and 24h, upon which an aliquot of cells was used to purify intact vacuoles. Total protein extract from complete protoplast and from vacuolar fraction were then subjected to trypsin treatment and 18-plex tandem mass tag (TMT) labelling, followed by HPLC-Orbitrap-MS analysis. **B.** Venn diagram illustrating the total number of proteins enriched in the vacuoles of shoots and roots upon 2h and 24h of autophagy induction. Consistent with our observations made in other experiments, root vacuoles accumulate proteins already after 2h of autophagy induction, while proteome of shoot vacuoles changes only after 24h of AZD/ConA treatment. Furthermore, in alignment of our previous observation, the number of proteins enriched in the root vacuoles during autophagy is significantly higher than in shoot vacuoles. The percentages represent the proportion of enriched proteins among the total number of proteins detected in the MS experiment for the specific organ. **C.** Diagram showing groups of selected hits enriched in the vacuoles of roots or shoot cells after 24h of autophagy induction. Names of individual interesting hits are shown by their respective groups. **D**. Venn diagram illustrating partial overlap of the proteins enriched in the vacuoles of shoot and roots cells upon 24h of autophagy induction. The percentages represent the proportion of hits that were enriched only in the specific organ but were detectable in the quantitative proteomes for both organs.

As our primary objective was to compare alterations in the vacuolar proteomes across a set of samples with naturally varying total protein concentration, we sought a reliable method to standardize our samples based on the amount of vacuoles they contain. We hypothesized that we could estimate the amount of vacuolar material originally present in each of our samples by normalizing the sample protein content to the detected amount of a vacuolar resident protein that is not impacted by the treatment. Good protein candidates for such normalization could be tonoplast-located aquaporins also known as intrinsic tonoplastic proteins (TIPs)^30^ or transmembrane subunits of the tonoplastic V-ATPase complex^31^. Therefore, we empirically verified that neither the localization nor the fluorescence intensity of the γTIP-GFP were affected by autophagy-inducing conditions (**Fig. S10E**), and the detectability of the protein by Western blot assay was also not impacted (**Fig. S10F**). Additionally, we confirmed that VHA-a3, one of the tonoplastic subunits of the V-ATPase complex, was also not detectable in the autophagic bodies upon induction of autophagy (**Fig. S10G)**. Therefore, we adhered to the strategy of normalizing detected proteomes to the corresponding detected amount of tonoplastic proteins before identifying differentially enriched hits.

Consistent with our previous findings of the earlier autophagic response in the roots, we observed protein accumulation in the root vacuoles already upon 2 hours of autophagy-inducing conditions, while in the shoot vacuoles it was detectable only after 24h of treatment (**Fig.5B, Tables S1** and **S2**). We detected a significantly larger number of proteins enriched in the root vacuoles upon 24h of autophagy induction (1307 proteins, 26% of total detectable root proteome **Fig.5B**) compared to the shoot vacuoles (506 proteins, 14 % of the total detectable shoot proteome **Fig.5B**) (**Tables S1-3**), further corroborating our observations of the stronger autophagic response in the roots.

Among the proteins that became enriched in the vacuoles upon induction of autophagy we could identify resident proteins of the organelles that we demonstrated to be sequestered into autophagosomes, such as mitochondria, Golgi apparatus, endoplasmic reticulum, and multivesicular bodies **(Fig.5C, Tables S1-3**). Consistent with our observations obtained using TEM and the fluorescent marker for cytoplasm uptake (mRFP), vacuolar proteomes of both plant organs exhibited a significant enrichment in cytoplasmic content, for example ribosomal proteins **(Fig.5C, Tables S1-3**). Intriguingly, we also observed that induction of autophagy caused enrichment of plastid or chloroplast proteins in the vacuolar proteomes of root and shoot cells **(Fig.5C, Tables S1-3**). To verify this finding, we induced autophagy in Arabidopsis seedlings expressing markers for plastid stroma (TKTP-GRX1-roGFP)^32^ or envelope (OE-YA)^33^. For this, seedlings were treated with AZD, –N and –C in the presence of ConA for 0-24h (**Fig.S11, Movies S2-4**). Indeed, we could observe accumulation of stromal plastid marker in the vacuole of both shoots and roots under AZD/ConA treatment (**Fig.S11, Movie S2**), thus corroborating the proteomics results. Noteworthy, both stromal and envelope markers were detectable in the vacuolar puncta of roots cells under all three stress conditions, indicating sequestration of the plastids by autophagosomes. However, the plastid envelope marker seemed to be excluded from the vacuolar puncta in the shoots of seedlings subjected to –N and –C (**Fig.S11, Movie S3** and **S4**), indicating potential selective exclusion of the plastid envelope proteins from degradation under these conditions and therefore a mechanism of chloroplast sequestration distinct from the one that takes place under AZD treatment.

The proteomics also revealed vacuolar enrichment of a number of proteins previously shown to contribute to or cross-talk with plant autophagy: components of the EXOCYST complex, components of the Ubiquitin Proteasomal System, PUX7/CDC48, Calcium-binding proteins, P-body proteins (**Fig.5C, Table S3**). Further comparison of proteins enriched in the root and shoot vacuole after 24h of autophagy induction revealed only partial overlap, thereby suggesting a significant organ-specific selectivity of AZD-induced autophagy (**Fig.5D**). The 177 proteins identified as common targets of autophagy in roots and shoots, mainly comprised ribosomal proteins and resident proteins of the organelles demonstrated in this study to be autophagic cargo in both plant organs (**Table S3**). Predictably, quantitative proteomes identified for shoots and roots were distinct, comprising 2326 proteins quantitatively detectable only in roots, 865 proteins unique for shoots and 2670 shared proteins. Notably, almost half (40%) of proteins enriched only in root vacuoles upon autophagy upregulation stemmed from the shared quantitative proteome, corroborating selectivity of AZD-induced autophagy. Similarly, more than half (62%) of proteins enriched only in shoot vacuole upon autophagy induction, were quantitatively detectable in the proteomes of both organs.

Remarkably, different sets of kinases involved in plant cell homeostasis were enriched in the root and shoot vacuoles of the same seedlings. Induction of autophagy in roots sequestered into the vacuoles MAPK1, MAPK4 and vps34, among which the latter is also directly involved in autophagy^34^; whereas shoot vacuoles were enriched for SnAK^35^ and STY46^36^ kinases regulating plant nutrient sensing, plant growth and abiotic stress responses (**Fig.5C, Tables S1-3**).

Intriguingly, we observed enrichment of phosphorylated ATG10 in the vacuoles of roots under prolonged stress (24h). ATG10 is an E2-like enzyme, a part of the two ubiquitin-like conjugation systems which are critical for autophagosome formation^37^. Other components of these systems were shown to undergo inhibitory phosphorylation by ATG1 during autophagosome maturation^38^, it is possible that plant ATG10 also undergoes phosphorylation by ATG1 and its subsequent uptake into the vacuole might be playing a role in dampening runaway autophagic activity under extended stress.

In summary, analysis of vacuolar proteomes fully supported our findings of autophagic activity occurring earlier and at a higher rate in roots compared to shoots. The proteomics data further confirmed uptake of cytoplasmic proteins, as we initially based on the results of mRFP marker abundance in vacuolar puncta and the presence of cytoplasm in the autophagic bodies on the TEM micrographs. Finally, vacuolar proteomes of shoot and root cells indicated strong organ-specific difference in autophagy cargo-selectivity.

## Discussion

To the best of our knowledge, this study is the first attempt to provide a comprehensive overview of the autophagic activity and selectivity simultaneously occurring in the shoots and roots under various stress conditions. By inducing autophagy using AZD8055, a selective ATP-competitive TORC1 inhibitor^15^, we observed the potential maximum of TOR-dependent autophagic activity and its selectivity in both plant organs. Further, comparison of these results with autophagy induced by physiologically relevant depletion of macronutrients, carbon or nitrogen, revealed the differences in the organ-specific approach to recycling its cell content depending on the stress (**Fig.6**).

**Figure 6.**
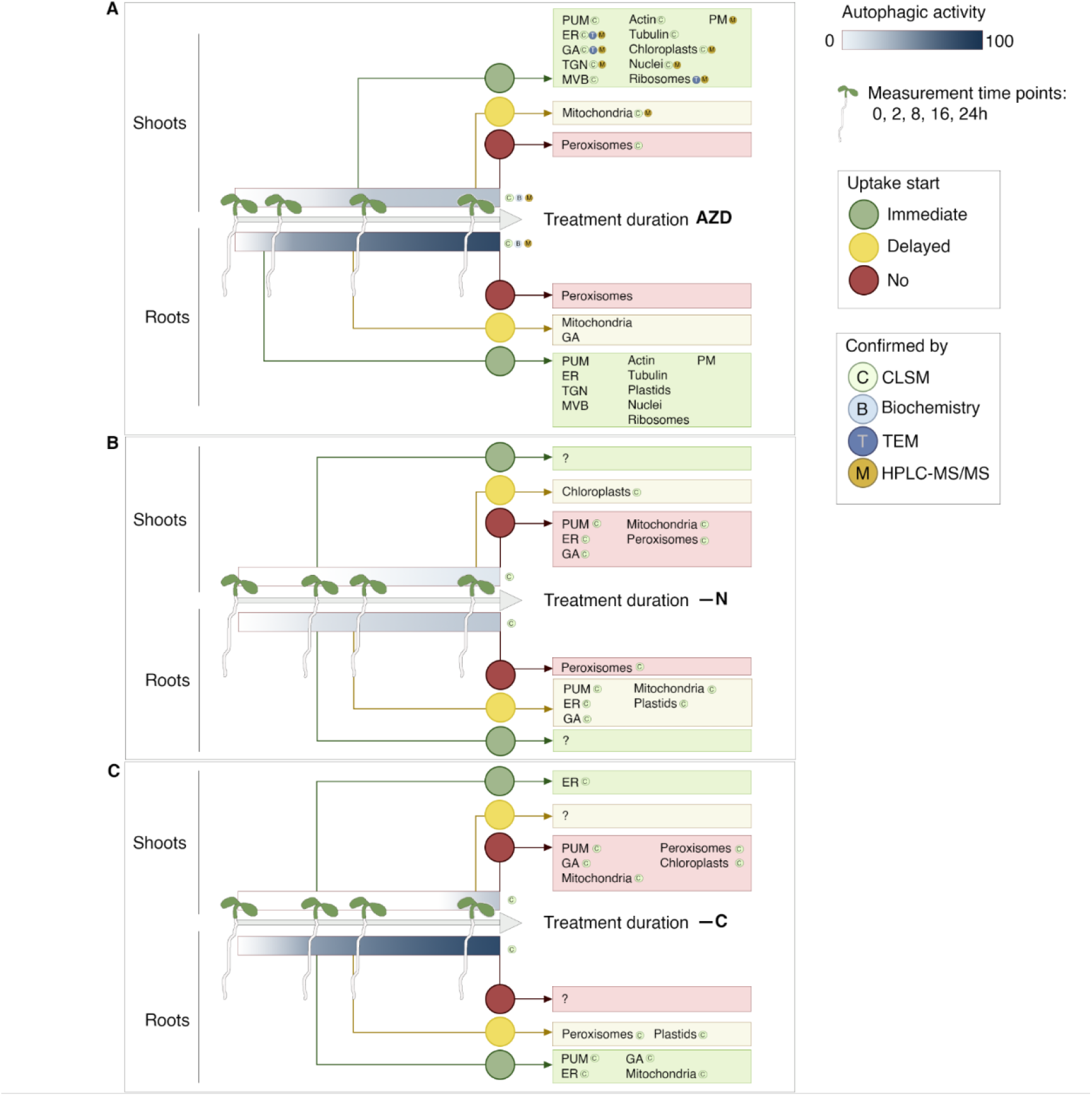
Spatio-temporal map of autophagic activity and its selectivity observed in Arabidopsis seedlings under three stress conditions. A diagram synthesizing results obtained in this study using a combination of cell biological and biochemical assays, electron microscopy, and proteomics. Under all three stress conditions, Arabidopsis roots consistently displayed higher activity and responded more rapidly than the shoots. AZD treatment triggered immediate sequestration of not only passive uptake marker (PUM) but also of various cell components in both shoots and roots. Notably, the uptake of mitochondria into autophagosomes was delayed in both organs under these conditions. Furthermore, GA sequestration was delayed in the roots only. Finally, pexophagy was not observed at all under treatment with AZD. These results indicate plant organ-specific selective protection of some cargo even under conditions that induce autophagy targeting a broad range of cell components, including passive cargo. Under nitrogen depleted conditions, the shoots exhibited strict selectivity of autophagy: only chloroplast markers were delivered to the vacuole; whereas in the roots upon prolonged treatment, autophagic cargo eventually encompassed a broader range cell components types. Similar to AZD-triggered autophagy, selectivity of –N-induced autophagy protected peroxisomes from being degraded in both plant organs. Intriguingly, –C conditions also yielded strict autophagic selectivity, which however, protected chloroplasts from degradation and instead targeted ER. Out of all checked conditions, pexophagy was observed only in the roots of seedlings subjected to prolonged carbon depletion. PUM, passive uptake marker (mRFP); ER, endoplasmic reticulum; GA, Goli apparatus; TGN, trans-Golgi network; MVB, multivesicular bodies; PM, plasma membrane

Initially, we hypothesized that autophagic activity might be elevated in the plant organ which nutrients are remobilized and reallocated to support the growth of another organ. Indeed, we observed that autophagy induced by carbon depletion showed much lower activity and more strict selectivity towards cell components in the shoots compared to roots (**Fig4C**, **Fig.S8**). It is plausible that roots activate autophagy at a higher rate and with a broader range of targets, because they become disproportionally more starved compared to the shoots that are preserving their local carbon supply to support etiolation.

However, the observations we made under nitrogen depletion showed us yet again that biology is never that predictable. Previous study showed that if environmental nitrogen is scarce, plants maintain higher levels of ammonium and amino acids in the roots at the expenses of the shoots ^8^. The higher nitrogen content under such conditions is vital to support TORC1 activity essential for promoting cell division and protein synthesis during foraging root growth. TORC1 is a well-described suppressor of autophagy^5,39^, therefore, we were surprised to discover a higher autophagic activity in the roots compared to shoots during –N treatments (**Fig.2G, Fig.S9**). One plausible explanation is that we detected a selective TORC1-independent autophagy, analogous to what has been previously demonstrated in animal models^11,40^. Furthermore, nitrogen-starved yeast cells were previously shown to selectively degrade the Ald6p enzyme, which was suggested to endanger cell viability under low nitrogen availability.^14^ TORC1-independent autophagy in animal cells is initiated by an autophagy receptor which recognizes the corresponding cargo and recruits the ATG machinery to it. Consequently, the autophagosomal membrane elongates in a tight proximity to the cargo surface, effectively preventing the unintended diffusion of other cargoes into the forming vesicle^11^. Our observations support this autophagosome formation strategy: in the beginning of –N treatment none of the autophagic bodies contained the reporter for soluble cytoplasmic proteins (mRFP, **Fig.4D** and **Fig.S9**), indicating strictly receptor-dependent target sequestration. The nature of the cargo within such selective plant autophagosomes remains and intriguing question for future studies for which our current work provides critical information on the timing of sampling. We demonstrate, that under prolonged –N treatment autophagy acquires a broader range of targets including cytoplasm, and certain organelles, with the exception of peroxisomes (**Fig.4D** and **Fig.S9**).

Furthermore, under –N conditions stromal marker of chloroplasts, but not its envelope marker was detectable in the vacuolar puncta (**Fig.S11**), indicating a selective exclusion of the chloroplasts’ envelope proteins during such chlorophagy. Notably, no chlorophagy was observed under –C conditions, albeit autophagic activity in the shoots was higher than under –N conditions (**Fig.5**, **Fig.S11**, **S8** and **S9**). It is tempting to speculate, that the most abundant plant protein RUBISCO located in the chloroplasts was used to replenish nitrogen source under –N and that chloroplasts were preserved under –C as a critical tool for the much-needed carbon assimilation in case if illumination conditions would become more favourable for photosynthesis.

Most intriguingly, we not only demonstrated that TORC1-dependent autophagy is significantly stronger and occurs quicker in the roots compared to shoots (**Fig.2**, **Fig.S7**) but also confirmed that this response pattern is intrinsic to the isolated cells of the corresponding organs (**Fig.S3**). What governs root cells propensity to a more rapid and robust autophagic response to TORC1 inhibition will be elucidated in future studies.

In this work, we also took a closer look into the broadly used and broadly debated concept of AZD-induced bulk autophagy^4,11^. Indeed, we observed that AZD induced formation of autophagic bodies containing varied mixes of sequestered cargoes (**Fig3.E**). However, we also demonstrated that peroxisomes were excluded from being sequestered into autophagosomes, while mitochondria and GA uptake into autophagosomes were delayed (**Fig.4D** and **C**), thereby demonstrating selectivity of autophagy induced by AZD. Therefore, AZD-induced autophagy likely comprises the co-occurring active and passive cargo sequestration strategies suggested in the **Fig.3D** and may not be accurately described as “bulk”.

Remarkably, we cold observe that all nine ATG8 isoforms of Arabidopsis localized to autophagic bodies upon AZD treatment of seedlings. Different ATG8s have been demonstrated to show distinct affinity towards known autophagy receptors.^41,42^ It would be interesting to test whether it is a combination of different Arabidopsis ATG8s decorating the same autophagosome that enables recruitment of multiple cargo types into the same vesicle under these conditions.

Most intriguingly, we also showed that while prolonged AZD-stress leads to an expected increase in autophagic activity in the shoots, it does not lead to an increase in the cytoplasmic marker uptake (mRFP) (**Fig.3B**). These results indicate that plant shoot autophagy cannot sustain targeting of a broad range of cell components for a prolonged period of time and eventually transitions to a stricter selectivity. Notably, no such decrease in the passive marker uptake was detectable in the root cells, showing their higher tolerance towards prolonged and massive degradation of the cellular content. It is possible that the intrinsic capacity of the root cells for the run-away autophagic activity is a survival strategy for the entire organism: under prolonged severe stress, the existing root sacrifices itself to support the shoot, which can form adventitious roots^43^ if later conditions become more favourable. Future studies will elucidate what prevents such response in the shoots and allows it in the roots.

We put a lot of effort into corroborating our conclusions implementing complementary approaches (**Fig.1**). However, more studies will be needed to provide a finer time-resolution and address unresolved questions. For example, –N and –C treatments had to be restricted to only 24h due to high cytotoxicity of these conditions in combination with ConA treatment. Development of non-invasive autophagy reporters, for example based on bioluminescence detection, would greatly aid further investigation of these stresses’ effects on plant autophagy.

For the sake of feasibility, in this study confocal microscopy on the roots was performed on the early differentiation zone, acquiring three technical replicates for each seedling (**Fig.S12**). However, we did not further dwell on the observed root zone-dependent differences in the autophagic activity, and recently reported only about the root cell-type dependent difference in the autophagic response to AZD/ConA treatment ^19^. Similarly, detection of autophagic activity in the leaves was performed using only adaxial side. We discovered that abaxial side of the same leaf exhibited little to no response to the treatment. This intriguing observation deserves further investigation in the future (**Fig.S12**).

In summary, this study presents the first time-resolved overview of autophagic activity occurring simultaneously in the photosynthetic autotrophic shoots and the heterotrophic roots of the same plant. We have shown that the timing, intensity, and selectivity of autophagic activity is plant organ-specific. These findings will serve as a roadmap for future research aimed at unraveling the intricate selectivity of plant autophagy.

## Acknowledgments

We would like to express our gratitude to Karin Staxäng and Monika Hodik (BioVis, Uppsala, Sweden) for their help with TEM experiments.

This study was supported by the funding from EU Horizon 2020 MSCA IF (799433 to EA Minina), Carl Tryggers Foundation (CTS 14 326 and 20:287 to EA Minina), The Swedish Research Council Formas (2021-01812 to EA Minina), Crops for the Future Research Program at the Swedish University of Agricultural Sciences and EPIC-XS (0000474 to EA Minina and S Holla).

## Materials and methods

### Plant material

Description of the stable transgenic lines used in this study is available in Supplementary table 5.

### Plant growth conditions

Arabidopsis seeds were surface sterilized in 10% bleach with 0.05% (v/v) Tween-20 for 30 minutes and washed thrice with sterile Milli-Q water. Surface sterilized seeds were plated on 0.5x Murashige and Skoog (MS) medium containing vitamins (M0222, Duchefa), supplemented with 10mM MES (pH 5.8) (M1503, Duchefa), 1% (w/v) sucrose, and 0.8% (w/v) plant agar (P1001, Duchefa). Seeds on plates were incubated vertically in a growth cabinet (Sanyo) under long-day growth conditions (16h of 120 µE m^−2^ s^−1^ light intensity at 22°C and 8h of dark at 20°C).

For experiments performed in RoPod v24, 3 mL of 0.5x MS medium was pipetted into sterile chambers. Approximately 0.3 cm of the agar was removed from the upper part of the chamber using a sterile scalpel. Sterilized seeds were then placed between the agar and the coverslip within each lane. The chambers were sealed and incubated horizontally under standard growth conditions for three days to allow the roots to reach the cover slip. Subsequently, the chambers were moved to a vertical position.

For plants grown in soil, 7-days-old seedlings grown on 0.5x MS medium were transferred to 8 cm^3^ pots filled with soil S-Jord (Hasselfors). Plants were grown under controlled growth conditions of 16 h light 8 h dark cycles, light intensity of 150 μE m^−2^ s^−1^ and 70% relative humidity.

### Sample preparation for confocal microscopy

To determine the most relevant time points for tandem-tag assay, RoPod v24 chambers ^19^containing 5-days-old seedlings were flooded with liquid 0.5x MS medium supplemented with 0.5 µM AZD8055 (AZD, Sigma Aldrich). The medium was replaced every 24 h with fresh 0.5x MS medium containing AZD, under sterile conditions. For the subsequent experiments, 5-days-old seedlings grown on 0.5x MS medium in Petri plates were transferred to six-well culture plates (3920300, Sarstedt) containing 3 mL 0.5x MS liquid medium supplemented with 0.5 µM AZD. The medium was replaced every 24 h with fresh 0.5x MS medium containing AZD.

For puncta quantification, 5-days-old seedlings grown on 0.5x MS medium were transferred to six-well plates containing 3 mL 0.5x MS liquid medium supplemented with 5 µM AZD8055 (364424, Santa Cruz Biotechnology) and 0.5 µM concanamycin A (ConA, 202111A, Santa Cruz Biotechnology). The plates were sealed and placed under standard growth conditions. The liquid medium containing AZD and ConA was replaced every 24h with fresh medium containing the drugs. For starvation-induced experiments, seedlings were transferred to carbon or nitrogen depleted medium supplemented with 0.5 µM ConA, and incubated under continuous dark or standard growth conditions, respectively.

For compound diffusion, 5-days-old seedlings grown on 0.5x MS medium were mounted in RoPod v23 chamber^19^ containing 0.5x MS liquid medium with 10 µg/mL Propidium iodide (PI) or 1 µg/mL Fluorescein Diacetate (FDA) or 1 µM FM4-64.

### Protoplast isolation from true leaves of Arabidopsis

Protoplasts were isolated from true leaves of 4-weeks-old GFP-ATG8 expressing plants, using the “Tape Arabidopsis Sandwich” described in Wu et al.^44^ The isolated protoplasts were collected in RoPod v22^19^ or 4-well glass bottom plates (P24-1.5H-N, Cellvis), and treated with 5 µM AZD and 0.5 µM ConA for the specified durations, before imaging.

### Protoplast extraction and vacuole isolation from Arabidopsis seedlings

Surface sterilized seeds were plated on Nunc Square BioAssay Dishes (240835, Thermo Fisher Scientific) containing 0.5x MS solid medium. Roots and shoots were harvested from 10-days-old seedlings and protoplasts were extracted from the two organs by using the method described in Wu et al ^44^. The roots and shoots were incubated in the enzyme solution for 16h in the dark, for efficient release of protoplasts. The extracted protoplasts were then treated with 5 µM AZD and 0.5 µM ConA for the specified durations. One-fourth of the protoplast fraction was frozen in liquid nitrogen. Intact vacuoles were isolated from the remaining protoplast fraction, using the method described in Robert et al.^45^, and was frozen in liquid nitrogen. The protoplast and vacuolar fraction were precipitated using 100% (w/v) Trichloro acetic acid (TCA). The experiments were performed to obtain three technical replicates, with each technical replicate consisting of a pool of three individual experiments.

### Confocal microscopy and data analysis

For tandem-tag assay and puncta quantification, seedlings were mounted in the corresponding liquid treatment medium, in RoPod v23 chambers.^19^ The root epidermal cells at the early differentiation zone were imaged using CLSM800 (Carl Zeiss), objective C-Apochromat 40×/1.2 W, excitation light 488 nm and 561 nm and emission ranges of (515–560 nm) and (570–650 nm) for GFP and TagRFP/mCherry, respectively. After imaging the roots, the cotyledons were excised, and mounted between coverslips in the corresponding treatment medium. The adaxial side of the cotyledons were imaged.

For compound diffusion, the root epidermal cells at the early differentiation zone and the adaxial side of the cotyledons were imaged using CLSM800 (Carl Zeiss), objective C-Apochromat 40×/1.2 W, excitation light 535 nm, 515 nm and 488 nm and emission ranges of (570–650 nm) and (515–560 nm) for PI, FM4-64 and FDA, respectively.

For imaging protoplasts isolated from true leaves, they were collected in RoPod v22 chambers or 24-well glass bottom plates and were imaged using CLSM800 (Carl Zeiss), objective C-Apochromat 40×/1.2 W, excitation light 488 nm and emission ranges of (515–560 nm).

Images were acquired using the Zen blue software (Carl Zeiss). For each experiment, all replicate images were acquired using the same confocal settings and the images were processed using ImageJ 1.53c.

For tandem-tag assay, the absolute integrated density in the green and red channels were measured using ImageJ 1.53c. The red to green ratio was calculated for each bio-replicate and the average of this was normalized to the average of the red to green values obtained from the autophagy deficient mutants.

Puncta quantification for the red and green channels was performed using the semi-automated image analysis ImageJ macro provided <u>here</u> (https://github.com/AlyonaMinina/Puncta-quantification-with-ImageJ). To quantify puncta accurately, extra caution was exercised to exclude cytoplasmic strands that could potentially resemble autophagic bodies **(Movie S5)**

### Transmission Electron Microscopy (TEM)

TEM was performed at BioVis, Uppsala. 5-days-old seedlings were treated with 5 µM AZD and 0.5 µM ConA, in 0.5x MS liquid medium. After treatment, the MS medium was replaced with a fixative containing 2.5% glutaraldehyde and 1% paraformaldehyde in 0.1 M phosphate buffer. The samples were post-fixed with 1% Osmium tetroxide and dehydrated in ethanol. The dehydrated samples were then embedded in resin and polymerized. Ultra-thin sections were made using an ultramicrotome. The sections were collected on copper slot and mesh grids and contrasted with uranyl acetate and lead citrate. After air drying, the sections were imaged using FEI Technai G2 operated at 80 kV. Images were processed using ImageJ 1.53c.

### Immunoblotting

For GFP-cleavage assay, surface sterilized seeds were plated on a 50-micron nylon mesh, placed on top of 0.5x MS medium. Plates were incubated vertically in the growth cabinet, under long day conditions. Plates containing 5-days-old seedlings was submerged in liquid 0.5x MS medium containing 5 µM AZD. For longer treatments, the liquid medium containing AZD was replaced every 24 h. For each time point, the roots and shoots were harvested and collected in separate Eppendorf tubes. The samples were ground in liquid nitrogen with a plastic pestle. The fine-powdered material was mixed with two volumes of 2x Laemmli buffer, thoroughly mixed, and boiled for 10 mins at 95°C. The samples were spun down at 13,000 g for 15 mins. The supernatant was loaded on Mini-PROTEAN TGX stain-free gels (4-20%; Bio-Rad), the proteins were transferred onto PVDF membranes, and stained with 1:1000 anti-GFP (632381, Clontech). The blots were developed using ECL Prime kit (RPN2232, Cytiva) and detected using Chemidoc (Bio-Rad). The intensities of the bands were quantified using ImageJ software. For each time point, the absolute integrated density was measured for bands corresponding to pHusion-ATG8 and free-GFP. The background values were subtracted from the absolute integrated density values. The sum of the values obtained for pHusion-ATG8 and free-GFP was set as 100%, and the free-GFP values were re-calculated as percentage of this.

### Liquid Chromatography and Mass Spectrometry

LC-MS/MS was performed at The VIB proteomics core, Gent, Belgium. Peptides from protoplast and vacuole samples of roots and shoots were isobarically labelled (Tandem Mass Tag, TMT-labelling) using 18-plex kit. The TMT labelled mixtures were fractionated on High-Performance Liquid Chromatography. The fractions were analyzed on the Orbitrap Fusion Lumos mass spectrometer.

The raw data from LC-MS/MS for all samples were searched together using the MaxQuant algorithm (version 2.2.0.0). The MS/MS spectra was searched against the Arabidopsis Uniprot database.

---

**Figure S1.**
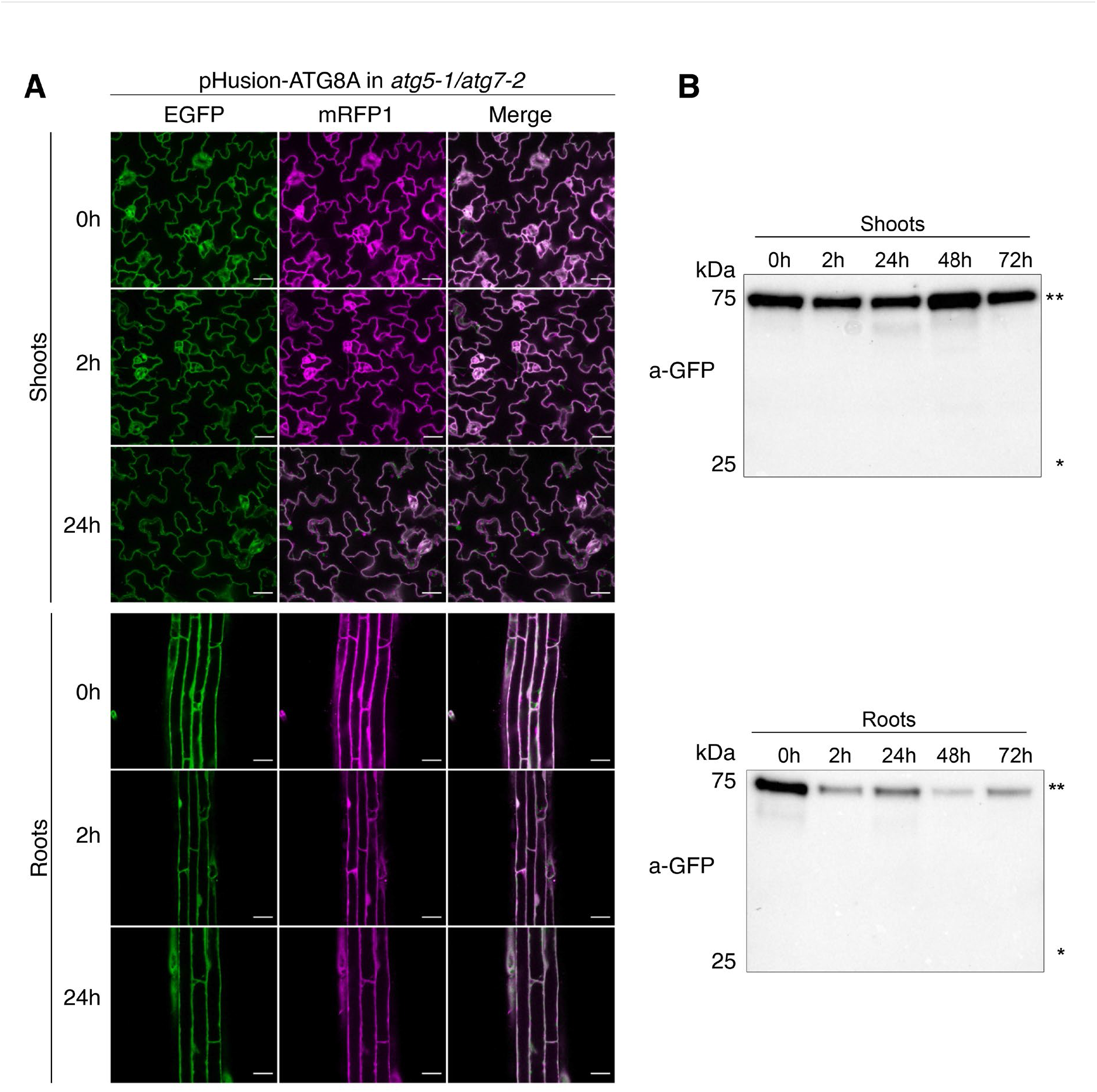
Delivery of pHusion-ATG8A to the vacuole is autophagy dependent. **A.** Confocal microscopy images obtained on the autophagy-deficient (*atg5-1/atg7-2)* seedlings expressing pHusion-ATG8A treated with AZD reveals no accumulation of the fluorescent marker in the vacuoles. The data shows the negative control obtained in the experiments illustrated in Fig.1A. Scale bars, 25 µm. **B.** GFP-cleavage assay shows only bands corresponding to the full length pHusion-ATG8 (**). *- represent the molecular weight corresponding to cleaved-GFP. The data shows the negative control obtained in the experiments illustrated in Fig.1C.

**Figure S2.**
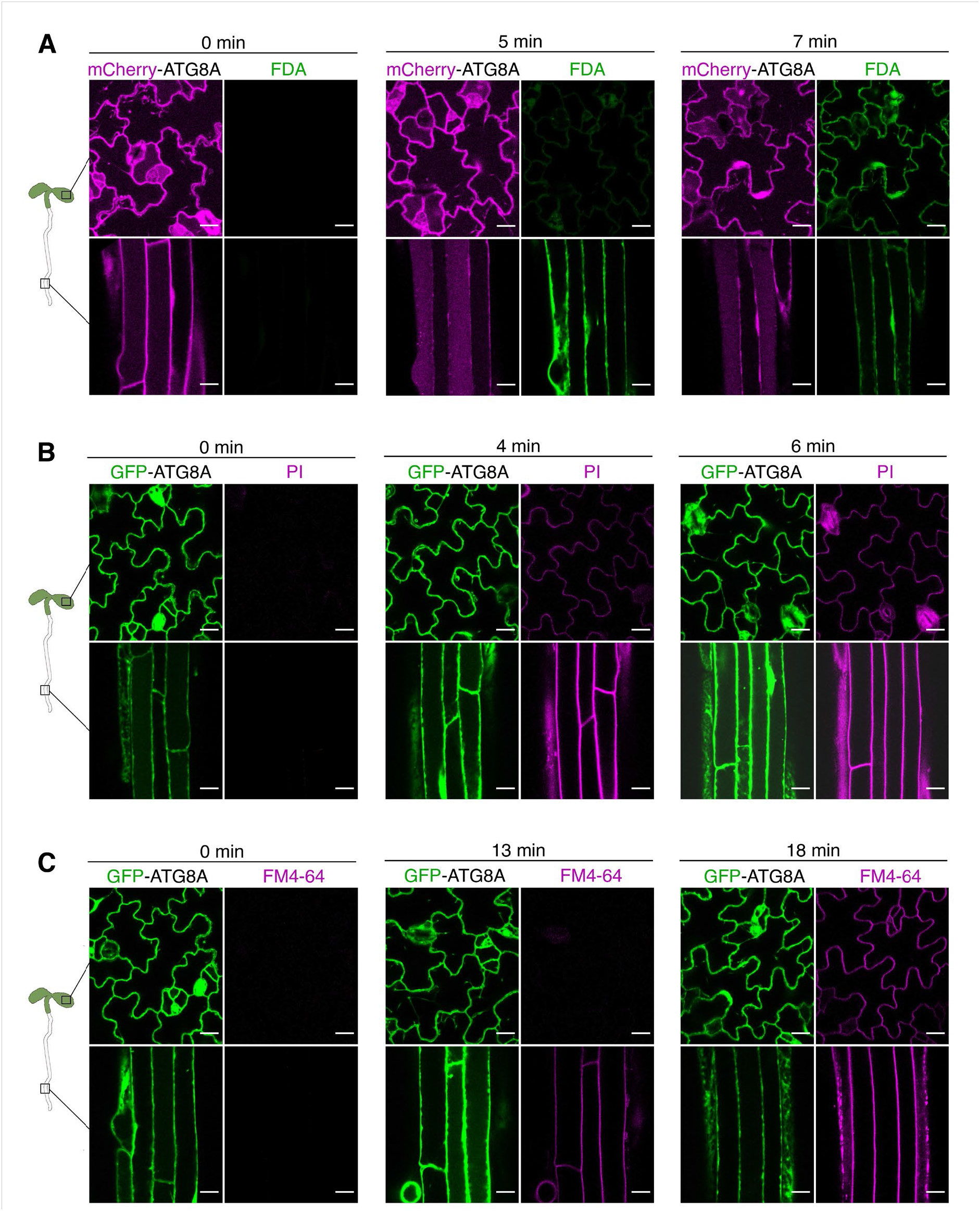
Shoots and roots of Arabidopsis seedlings exhibit comparable cell permeability. Cell permeability in the intact organs of Arabidopsis seedling was assessed by detecting time-resolved accumulation of small fluorescent organic molecules using CLSM. **A.** Uptake of Fluorescein diacetate (FDA) in both shoots and roots of mCherry-ATG8A-expressing seedlings occurs under ten minutes. Similarly, no major organ-specific difference in the uptake time for Propidium iodide (**B**) or FM4-64 (**C**) was observed in GFP-ATG8 seedlings. Scale bars, 15 µm.

**Figure S3.**
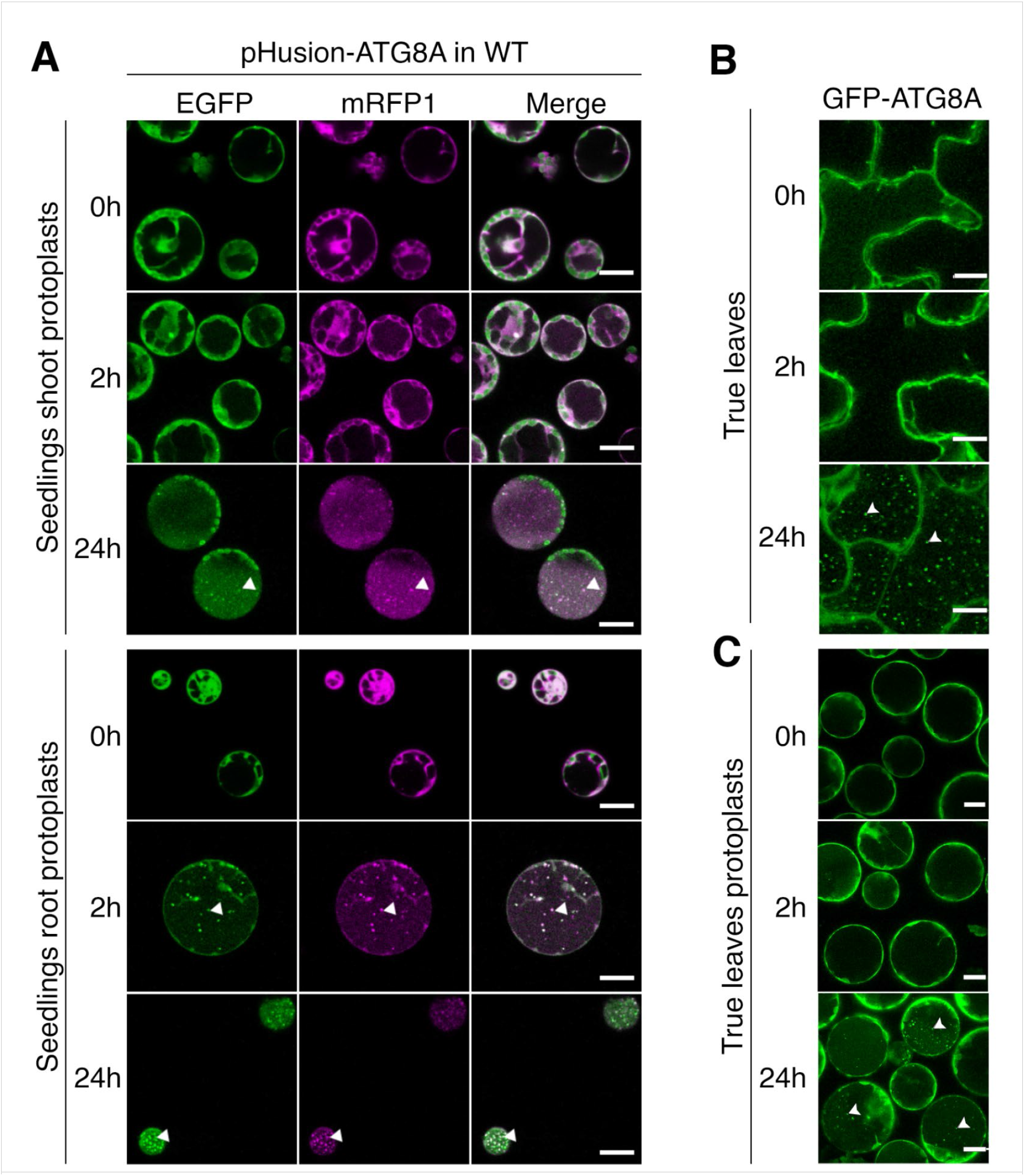
The organ-specific dynamics of autophagy is reproducibly observed in shoot and root cells isolated from seedlings and in true leaves. **A.** Confocal micrographs of protoplasts isolated from the shoots (top panels) and roots (bottom panels) of the seedlings expressing pHusion-ATG8A in the WT background. Isolated cells were treated with AZD/ConA for 0h, 2h and 24h prior to imaging. In agreement with observations obtained on the corresponding intact organs, autophagic activity is detectable in the isolated root cells already after 2h of treatment and in isolated shoot cells only after 24h. Scale bars, 20 µm. White arrowheads point at autophagic bodies in the vacuoles. **B.** Confocal microscopy performed on true leaves of 5-week-old plants expressing GFP-ATG8A. Leaves were infiltrated with AZD/ConA and scanned after 0h, 2h and 24 hours of treatment. Scale bar, 10 µm. White arrowheads point at autophagic bodies in the vacuoles. Protoplasts extracted from true leaves of GFP-ATG8A-expressing plants were treated with AZD/ConA 0h, 2h and 24 hours prior to imaging. Scale bar, 30 µm. White arrowheads point at autophagic bodies in the vacuoles.

**Figure S4.**
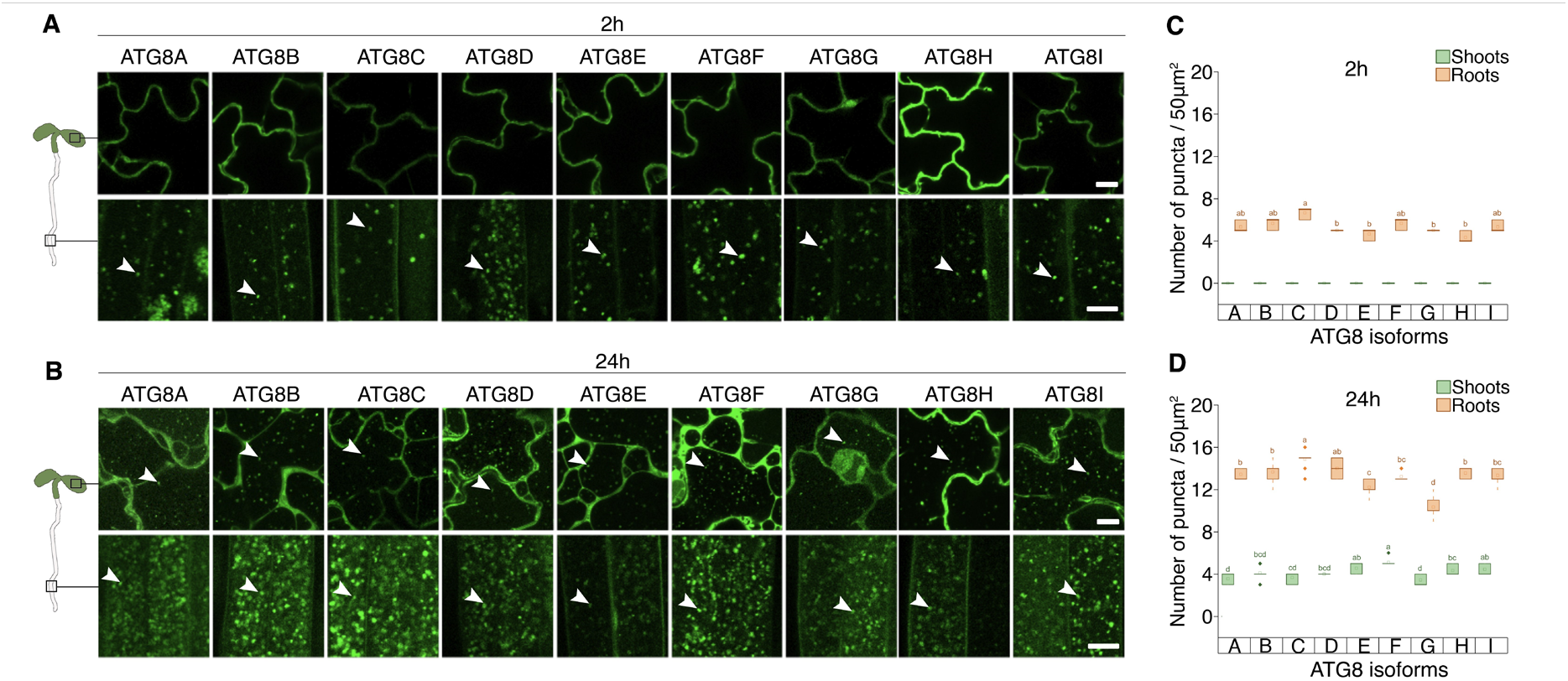
Earlier and stronger autophagic response in roots is reproducibly detectable with all nine ATG8 isoforms of Arabidopsis. **A** and **B.** Confocal microscopy performed on 5-days-old Arabidopsis seedlings expressing GFP-tagged ATG8 isoforms (GFP-ATG8A-I). Seedlings were treated with AZD/ConA for 2h (**A**) and 24h (**B**) prior to imaging. White arrowheads indicate autophagic bodies in the vacuoles. Scale bars, 10 μm. **C** and **D.** Quantification of autophagic bodies illustrated in (**A** and **B**) reveals higher autophagic activity in the roots compared to the shoots detectable with all nine ATG8 isoforms. The charts comprise data from a single representative experiment. Tukey’s test, groups statistically not different from each other are annotated by the same letter, *significance level 0.05*. Median is indicated by a line across the box; mean is indicated by small squares; range within 1.5IQR is represented by whiskers; outliers are represented by solid rhombus.

**Figure S5.**
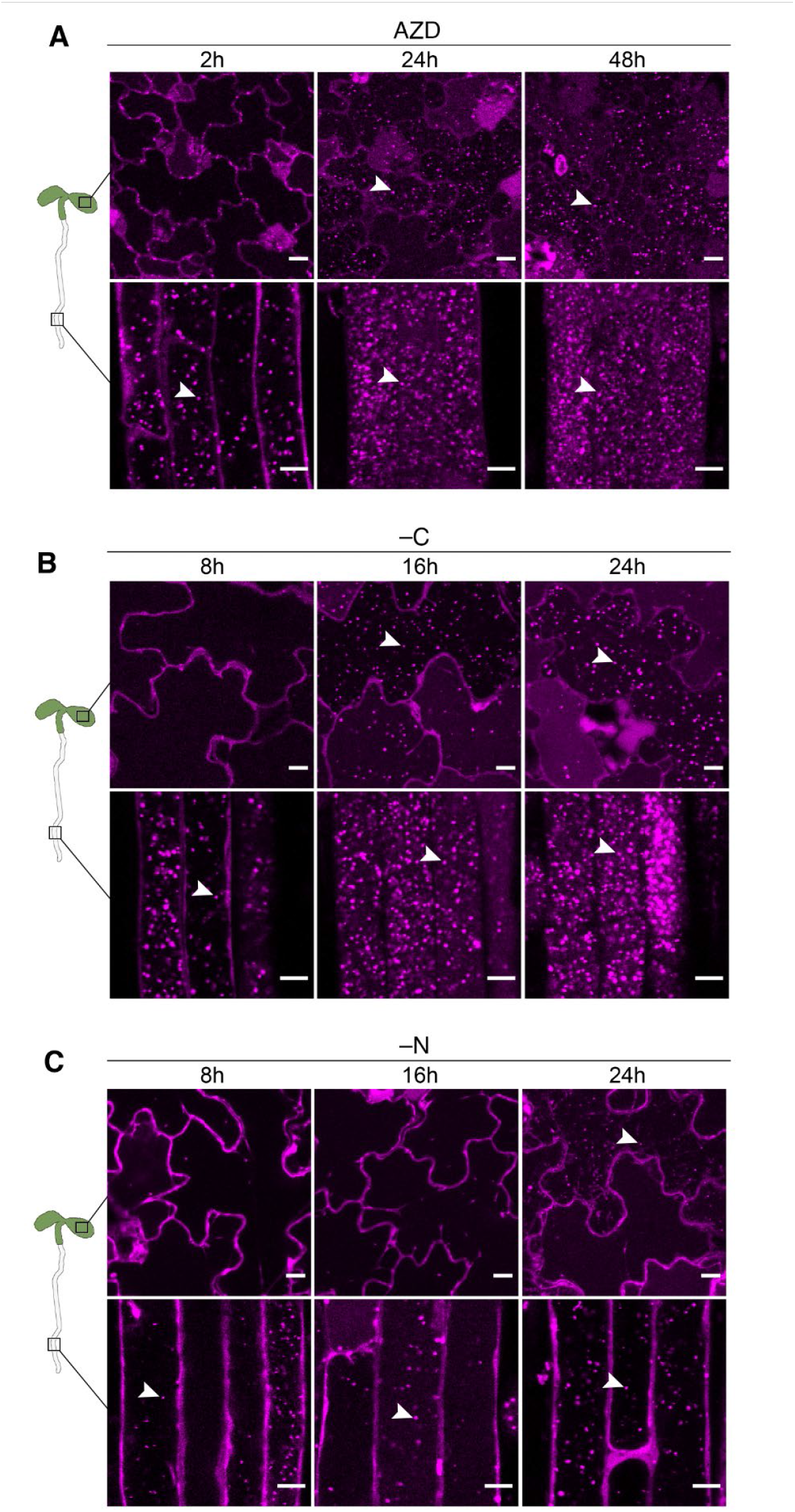
Earlier and stronger autophagic response in roots is reproducibly detectable with all three stress triggers. Confocal microscopy images obtained on 5-days-old Arabidopsis seedlings expressing mCherry-ATG8A. Seedlings were subjected to AZD treatment **(A)**, carbon depletion **(B)**, or nitrogen depletion (**C**). All treatments were carried out in the presence of ConA to prevent degradation of autophagic bodies. White arrowheads indicate autophagic bodies in the vacuoles. Scale bars, 20 µm.

**Figure S6.**
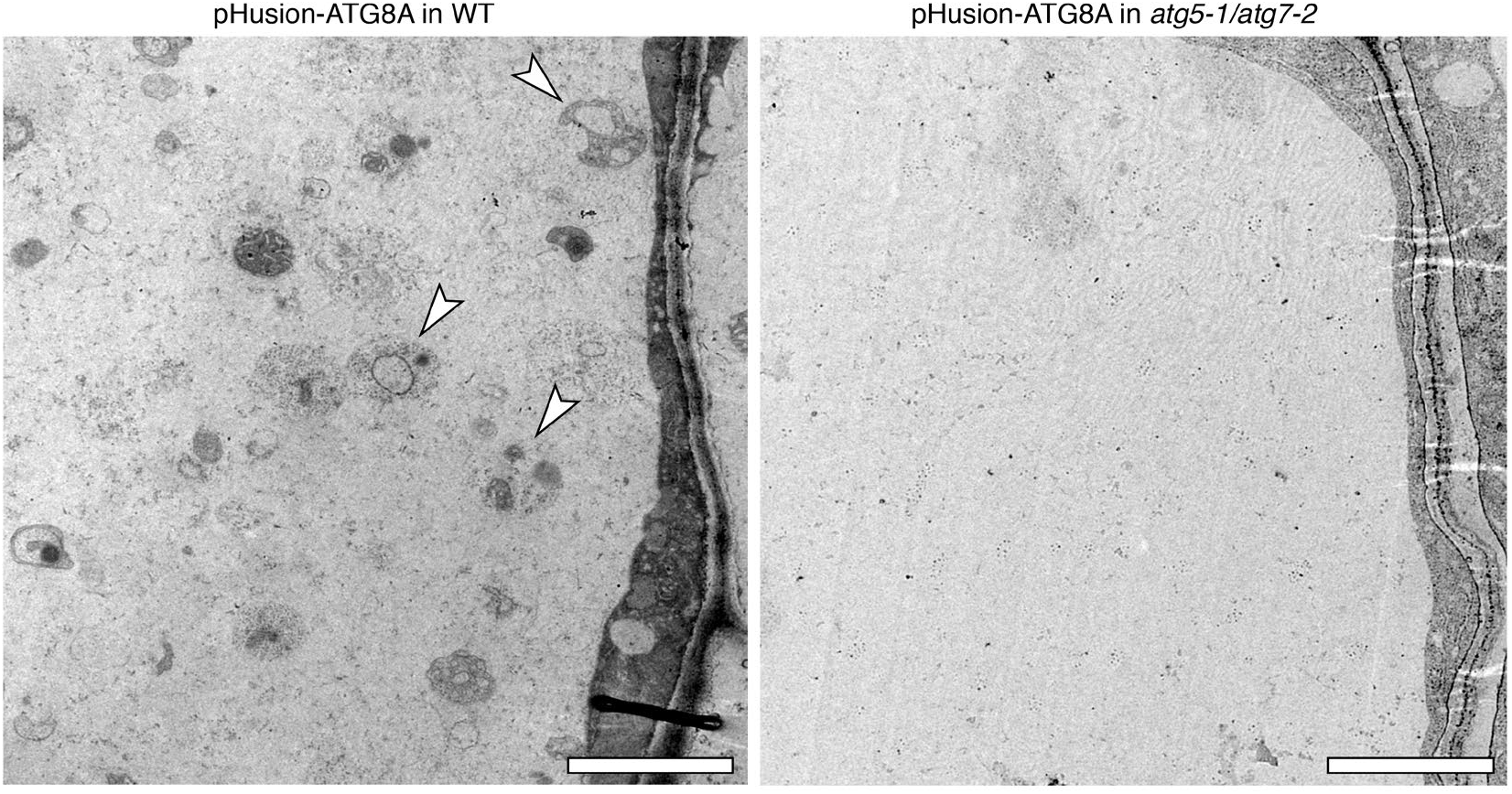
Structures annotated as autophagic bodies in the WT cells are absent from the vacuole of atg5-1/atg7-2 KO cells. Transmission Electron Microscopy (TEM) images obtained on the roots of seedlings expressing pHusion-ATG8 in the WT and autophagy-deficient (atg5-a/atg7-2) backgrounds. Seedlings were treated with AZD/ConA for 24h prior to sampling for TEM. White arrowheads point at autophagic bodies accumulated in the vacuole of the WT cell. Scale bars, 2 µm.

**Figure S7.**
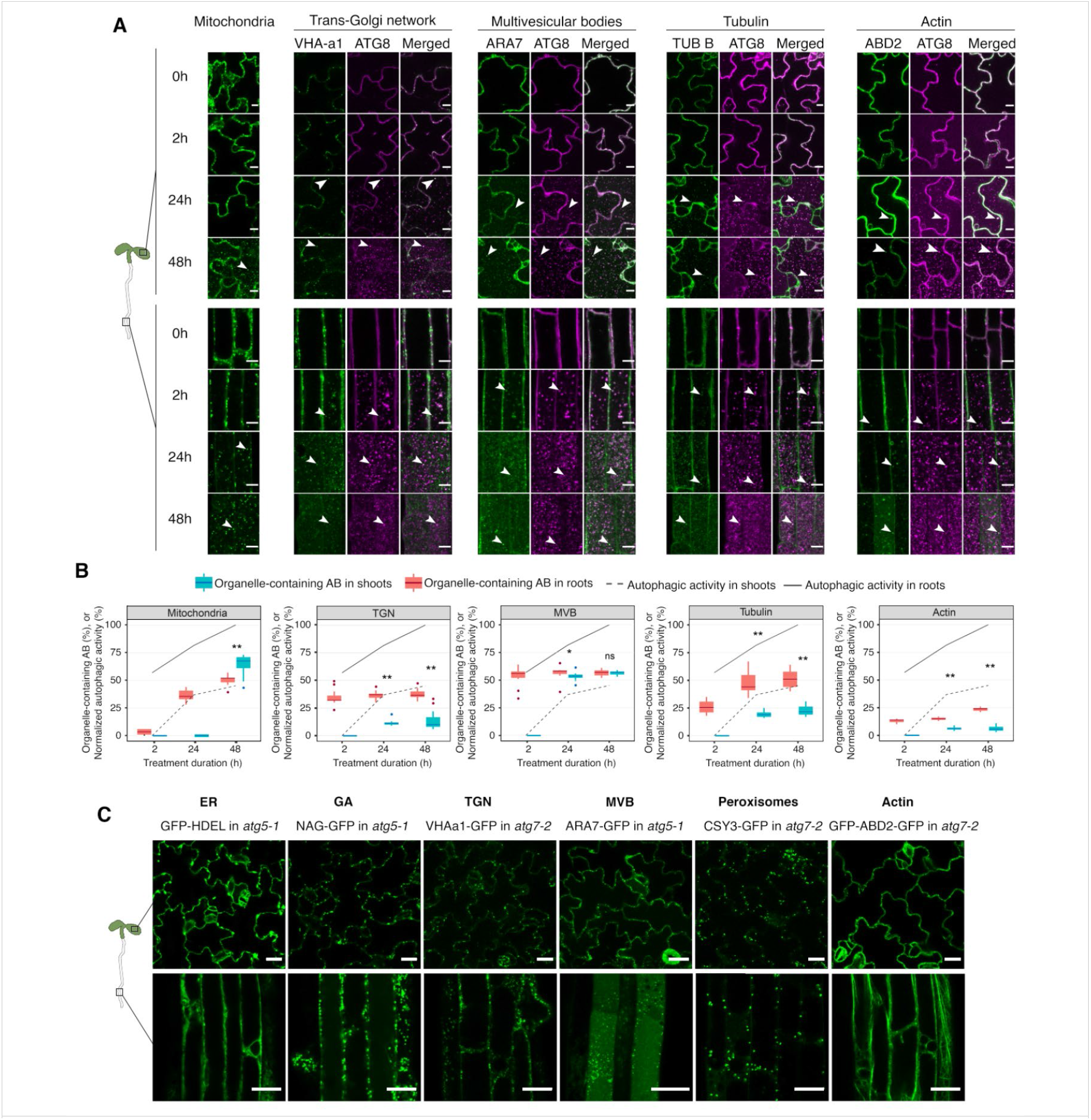
AZD-induced autophagy displays distinct spatiotemporal specificity towards organelles. **A.** Confocal micrographs obtained on 5-days-old Arabidopsis seedlings co-expressing mCherry-ATG8 with one of the following organelle markers: VHA-a1-GFP (trans-Golgi network, *TGN*), ARA7-GFP (multivesicular bodies, *MVBs*), Tub B-GFP (tubulin and microtubules), GFP-ABD2-GFP (actin). Seedlings expressing Mt-YFP (mitochondria) were implemented for tracking mitophagy. Seedlings were treated with AZD/ConA for the indicated amount of time. Filled arrowheads indicate autophagic bodies containing an organellar marker, outlined arrowheads point at autophagic bodies without the corresponding organellar marker. Scale bars, 10 µm. **B.** The charts represent the autophagic activity detected in the shoots (dotted line) and in the roots (solid lines), along with the percentage of autophagic bodies containing the specific organelle (box plots). Data represents two independent experiments. n=72, unpaired two-tailed Student’s t-test; *, p-value<0.05; **, p-value<0.0001; ns, not significant. **C.** Confocal micrographs illustrating that delivery of organelles to the vacuole upon AZD treatment is autophagy-dependent. Five-days-old Arabidopsis seedlings expressing the indicated organelle marker in autophagy-deficient background were subjected to 24h of AZD/ConA treatment. None of the organellar markers was delivered to the vacuole upon treatment, except for MVB marker which was detectable as diffuse signal in the vacuolar lumen. The latter could result from normal MVB fusion with the vacuoles, which occurs during anterograde endomembrane trafficking. Scale bars, 25 µm.

**Figure S8.**
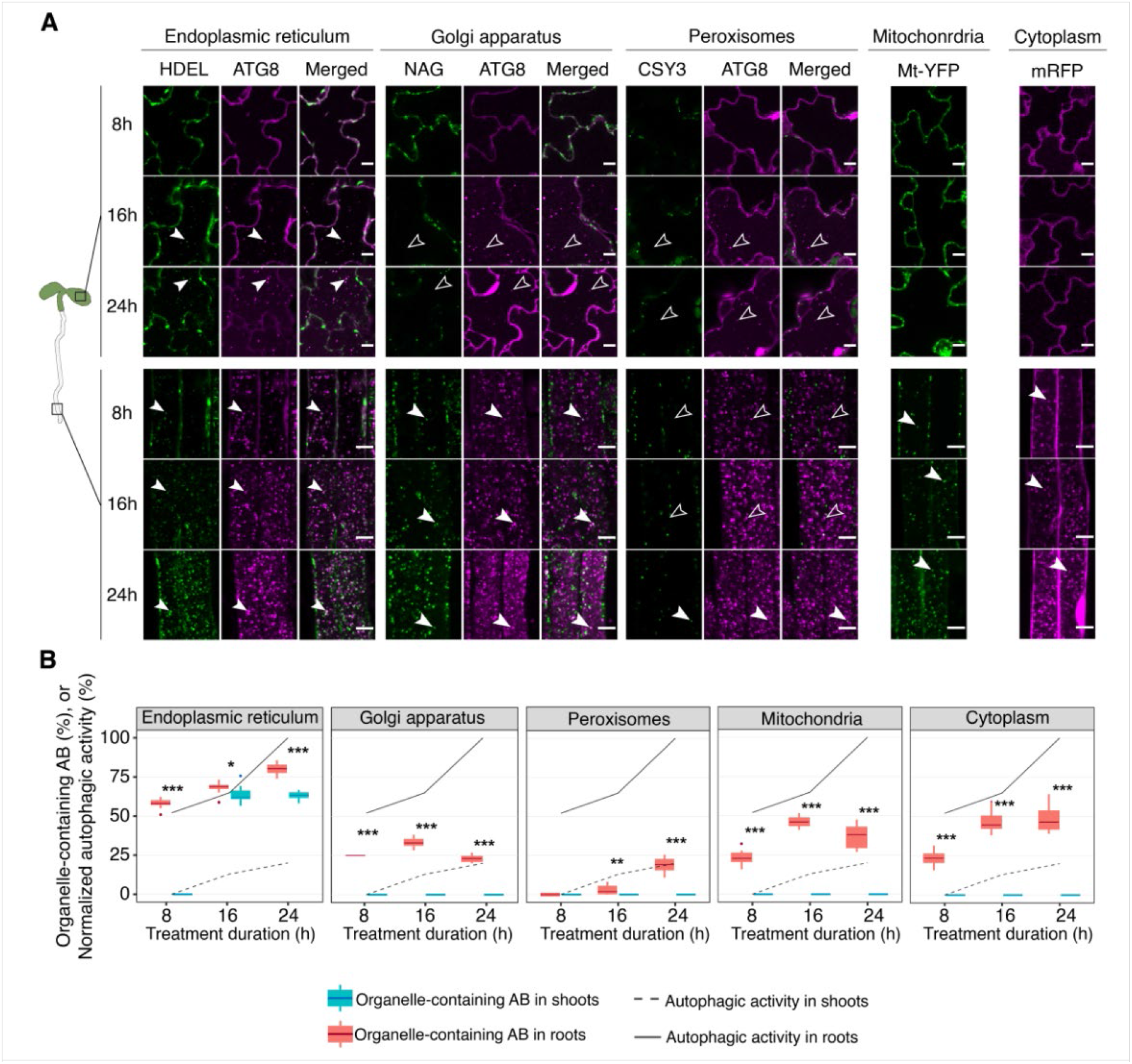
Autophagy displays distinct spatiotemporal specificity towards organelles under carbon depleted conditions. **A.** Confocal micrographs obtained on 5-days-old Arabidopsis seedlings co-expressing mCherry-ATG8 with one of the following organelle markers: GFP-HDEL (endoplasmic reticulum), NAG-GFP (Golgi apparatus) and CSY3-GFP (peroxisomes). Seedlings expressing Mt-YFP (mitochondria) were implemented for tracking mitophagy, while seedlings expressing mRFP were used as a marker for passive cargo uptake by autophagy. Seedlings were subjected to carbon depletion and simultaneously treated with ConA for the indicated amount of time. Filled arrowheads indicate autophagic bodies containing an organellar marker, outlined arrowheads point at autophagic bodies without the corresponding organellar marker. Scale bars, 10 µm. **B.** The charts represent the autophagic activity detected in the shoots (dotted line) and in the roots (solid lines), along with the percentage of autophagic bodies containing the specific organelle (box plots). Data represents two independent experiments. n=64, unpaired two-tailed Student’s t-test; *, p-value<0.01; **, p-value<0.001; ***, p-value<0.0001; ns, not significant.

**Figure S9.**
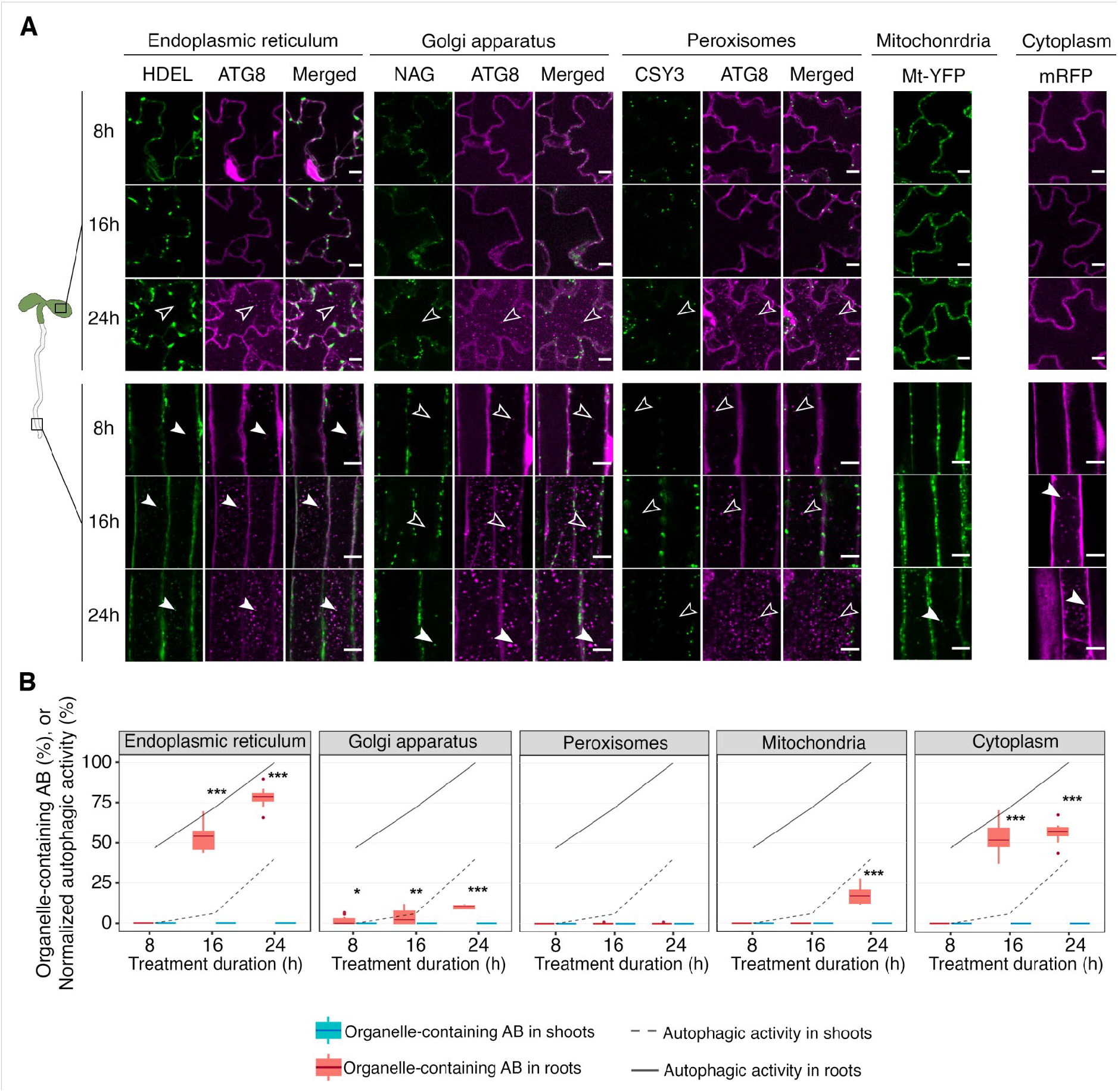
Autophagy displays distinct spatiotemporal specificity towards organelles under nitrogen depleted conditions. **A.** Confocal micrographs obtained on 5-days-old Arabidopsis seedlings co-expressing mCherry-ATG8 with one of the following organelle markers: GFP-HDEL (endoplasmic reticulum), NAG-GFP (Golgi apparatus) and CSY3-GFP (peroxisomes). Seedlings expressing Mt-YFP (mitochondria) were implemented for tracking mitophagy, while seedlings expressing mRFP were used as a marker for passive cargo uptake by autophagy. Seedlings were subjected to carbon depletion and simultaneously treated with ConA for the indicated amount of time. Filled arrowheads indicate autophagic bodies containing an organellar marker, outlined arrowheads point at autophagic bodies without the corresponding organellar marker. Scale bars, 10 µm. **B.** The charts represent the autophagic activity detected in the shoots (dotted line) and in the roots (solid lines), along with the percentage of autophagic bodies containing the specific organelle (box plots). Data represents two independent experiments. n=56, unpaired two-tailed Student’s t-test; *, p-value<0.05; **, p-value<0.01; ***, p-value<0.001; ns, not significant.

**Figure S10.**
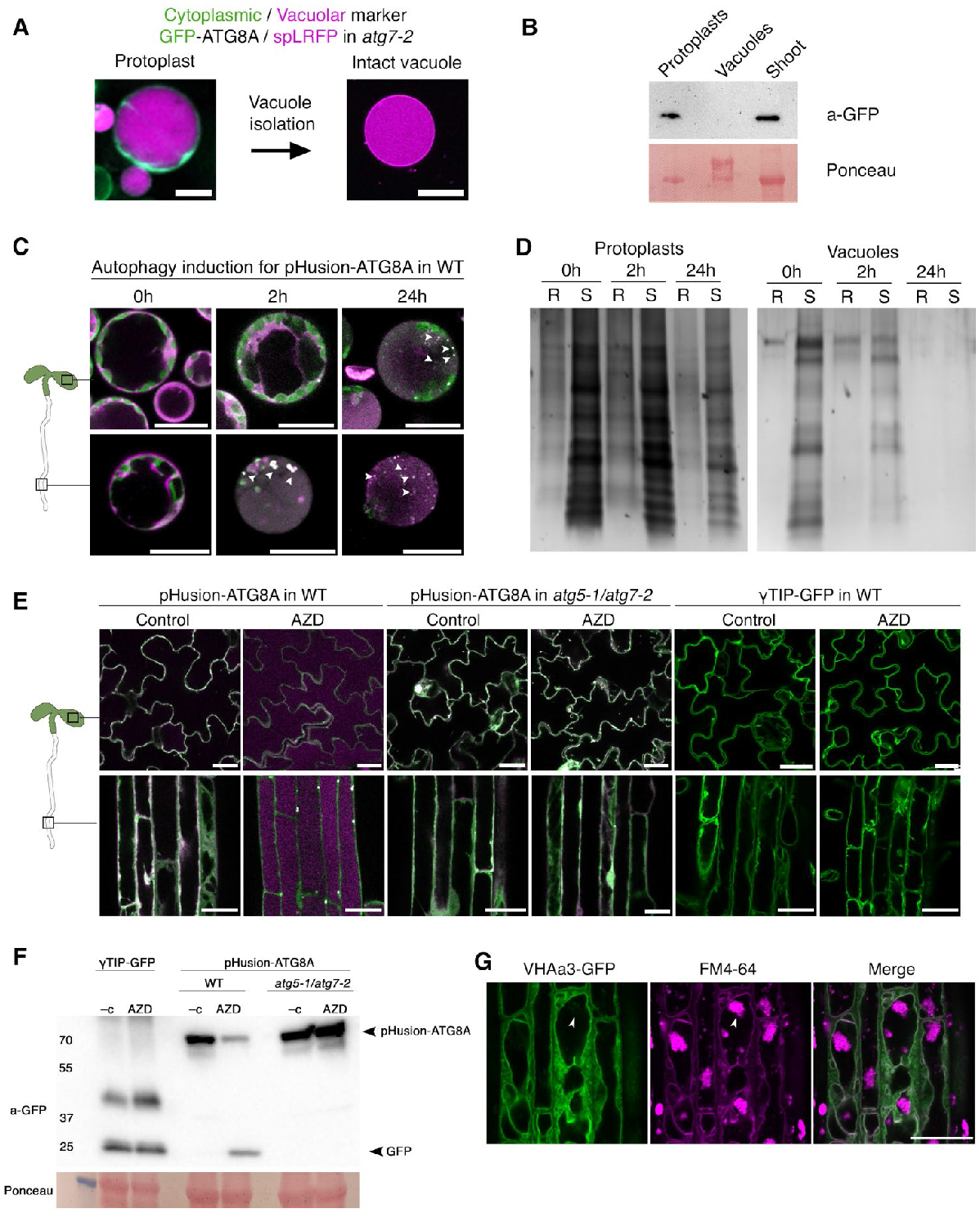
Verification of the strategy for proteomics experiments. **A.** Isolation of intact lytic vacuoles from shoots and roots of *Arabidopsis thaliana* plants co-expressing red vacuolar marker (SpL:RFP) and green cytoplasmic marker (GFP-ATG8) in the *atg7-2*. In the autophagy-deficient *atg7-2* background GFP-ATG8 remains in the cytoplasm even under autophagy-inducing conditions. Micrograph of the intact protoplast (left) and isolated vacuole (right). Scale bar, 25μm. **B.** Western blot performed on the samples shown in **A**. Lack of GFP signal in the vacuolar fraction confirms that isolated intact vacuoles are not contaminated with cytoplasmic proteins. Ponceau staining was used as a loading control. **C.** Confocal microscopy of protoplasts isolated from shoots and roots of WT plants expressing pHusion-ATG8 and treated AZD/ConA for the indicated amount of time. Accumulation of autophagic bodies in the vacuole confirmed autophagic response previously observed in the intact plant organs. Scale bar, 25μm. **D.** Silver staining of an SDS-PAGE gel showing total protein extracts obtained from protoplasts and from intact vacuoles. Protoplast were isolate from shoots (S) and roots (R), and treated with AZD/ConA for 0h, 2h and 24 hours prior to protein extraction. **E.** Confocal micrographs of cotyledons and roots from seedlings under control conditions and upon 24h of treatment with 5 μM AZD. Seedlings expressing pHusion-ATG8 in WT and autophagy-deficient background were used as a control for the efficacy of drug treatment, while seedlings expressing γTIP-GFP were used to check the effect of autophagy on the tonoplastic marker. Microscopy shows autophagy-dependent accumulation of the pHusion-ATG8 signal in the vacuoles of WT cells in both plant organs and demonstrates no discernible effect of autophagy induction on the localization or intensity of γTIP-GFP. Scale bar, 25μm. **F.** Western blot assay performed on the protein extract from the seedlings illustrated in **E**. Accumulation of the cleaved-off GFP in the WT background corroborates efficient upregulation of autophagy under used conditions. γTIP-GFP signal appears unaffected under the same conditions. **G.** Confocal micrographs of seedlings roots expressing tonoplastic marker VHAa3-GFP and treated with AZD/ConA for 24h. Seedlings were pretreated with FM4-64 prior to induction of autophagy to visualize autophagic bodies accumulating in the vacuolar lumen. Microscopy revealed no uptake of the tonoplastic marker upon autophagy induction. Scale bar, 25μm.

**Figure S11.**
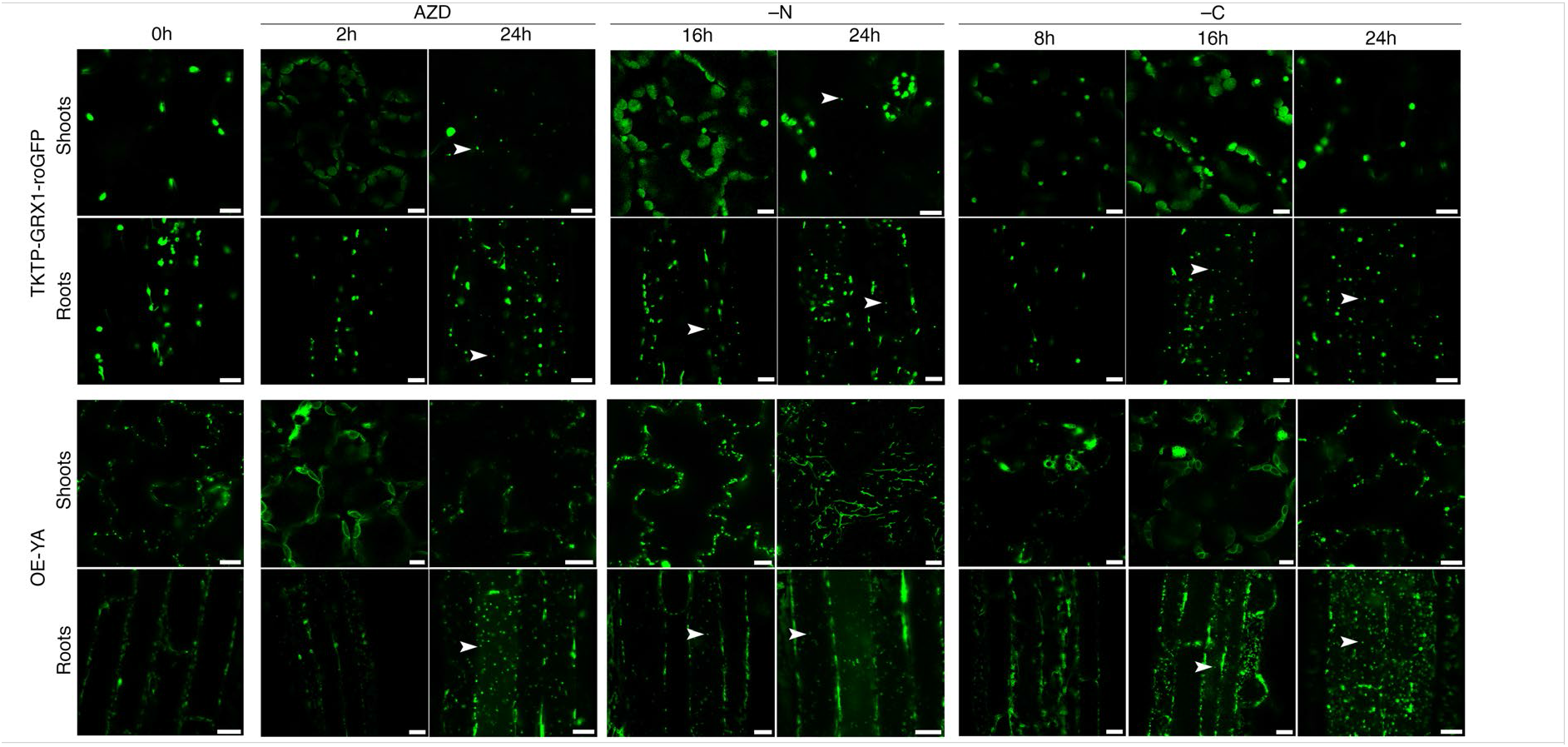
Translocation of plastid markers in the vacuoles of seedlings upon autophagy induction. Confocal micrographs obtained on Arabidopsis seedlings expressing stromal marker (TKTP-GRX1-roGFP) or envelope marker (OE-YA) for plastids/chloroplasts and subjected to autophagy induction by treatment with AZD, – N or – C for the indicated amount of time. All treatments were carried out in the presence of ConA to prevent degradation of a marker in the lytic vacuoles. Scale bar, 10 μm. White arrowheads point out at the vacuolar puncta.

**Figure S12.**
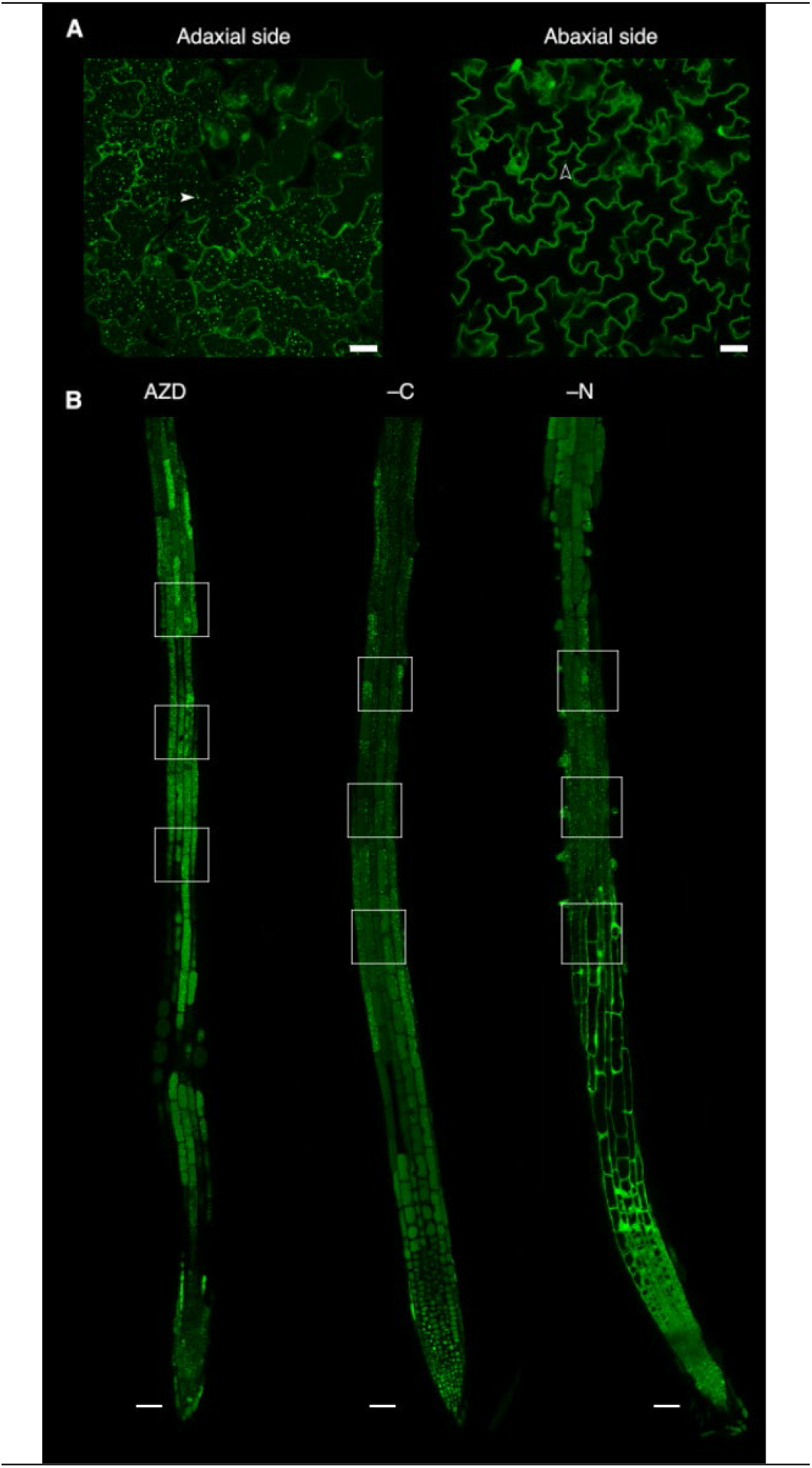
Abaxial and adaxial leaf sides as well as different root zones show significant variation in the accumulation of autophagic bodies. **A.** An illustrative confocal micrographs of the abaxial and adaxial sides of a cotyledon expressing GFP-ATG8A in the WT background. The seedling was treated with AZD/ConA for 24h prior to imaging. Filled white arrowhead indicates an autophagic body, outlined arrowheads indicates a cytoplasmic strand. The low number of autophagic bodies in the abaxial epidermal cells was consistently observed in four independent experiments using three different markers for autophagosomes. In this study only adaxial side of true leaves and cotyledons was used for imaging. Scale bars, 25 µm. **B.** Tile scan images of 5-day-old roots expressing GFP-ATG8A. Seedlings were exposed to 24h treatment with AZD, –C (carbon depletion) or –N (nitrogen depletion) in the presence of ConA. To ensure comparability of the results in this study, confocal microscopy of roots under all treatments was performed within the region spanning from the late elongation to the early differentiation zones, three images (technical replicates) were acquired for each seedling (biological replicate). Approximate positions of scans are indicated by the white rectangles. Scale bars, 50 µm.

**Movie S1.**
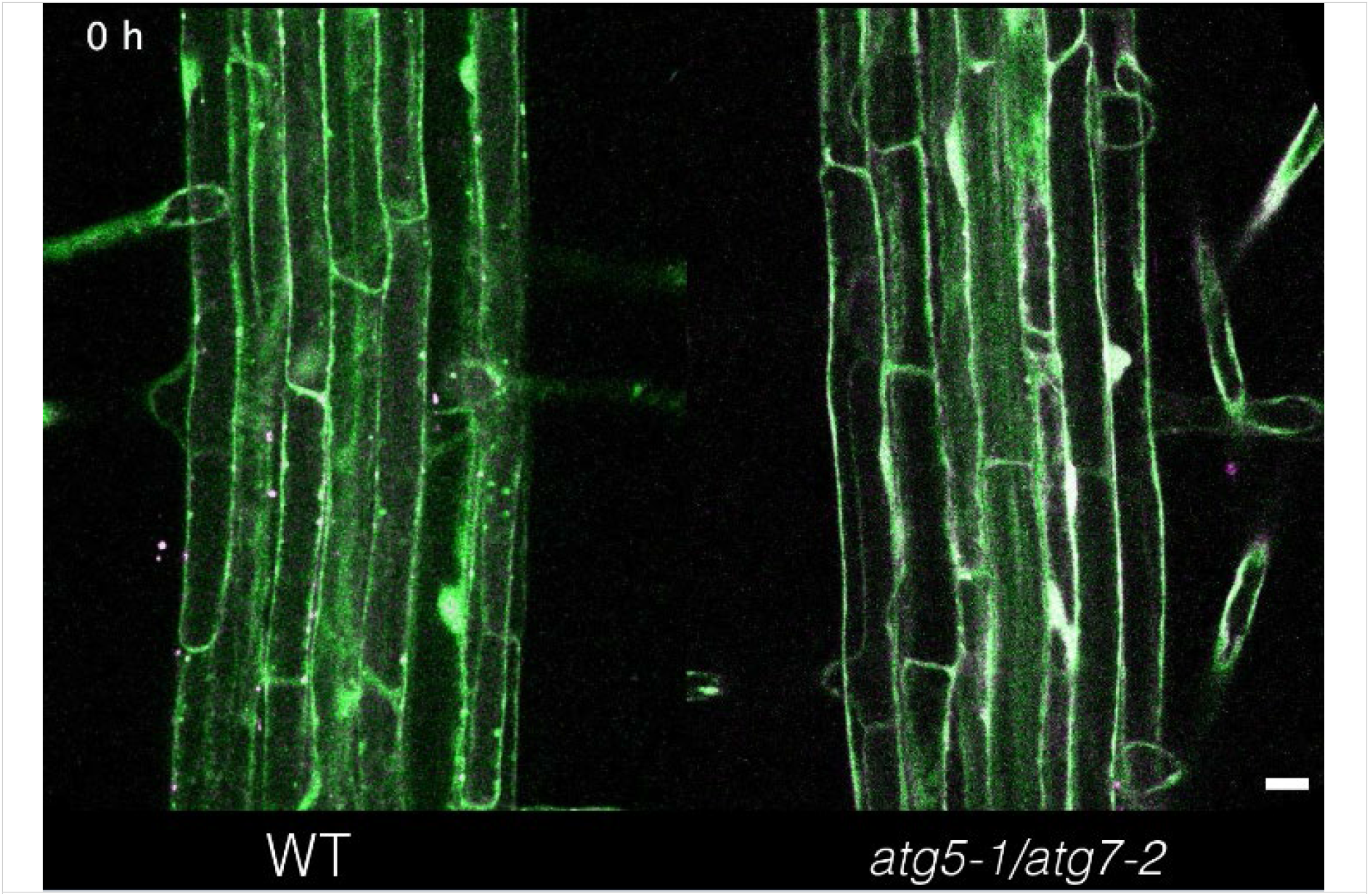
Time-resolved detection of autophagic activity in *Arabidopsis thaliana* roots. Arabidopsis seedlings expressing pHusion-ATG8A (EGFP-mRFP1-ATG8A) in the WT (left) and in the autophagy-deficient *atg5-1/atg7-2* (right) backgrounds were grown in the RoPod chamber (Guichard et al., 2021). AZD8055 was added to the growth medium within the chamber, and seedlings were imaged every hour for 3 days. Fluorescent marker is detectable in the WT root vacuoles already upon 2 hours of treatment, whereas the signal remained cytoplasmic in the autophagy deficient mutants for the whole duration of the experiment. Scale bar, 50 µM.

**Movie S2.**
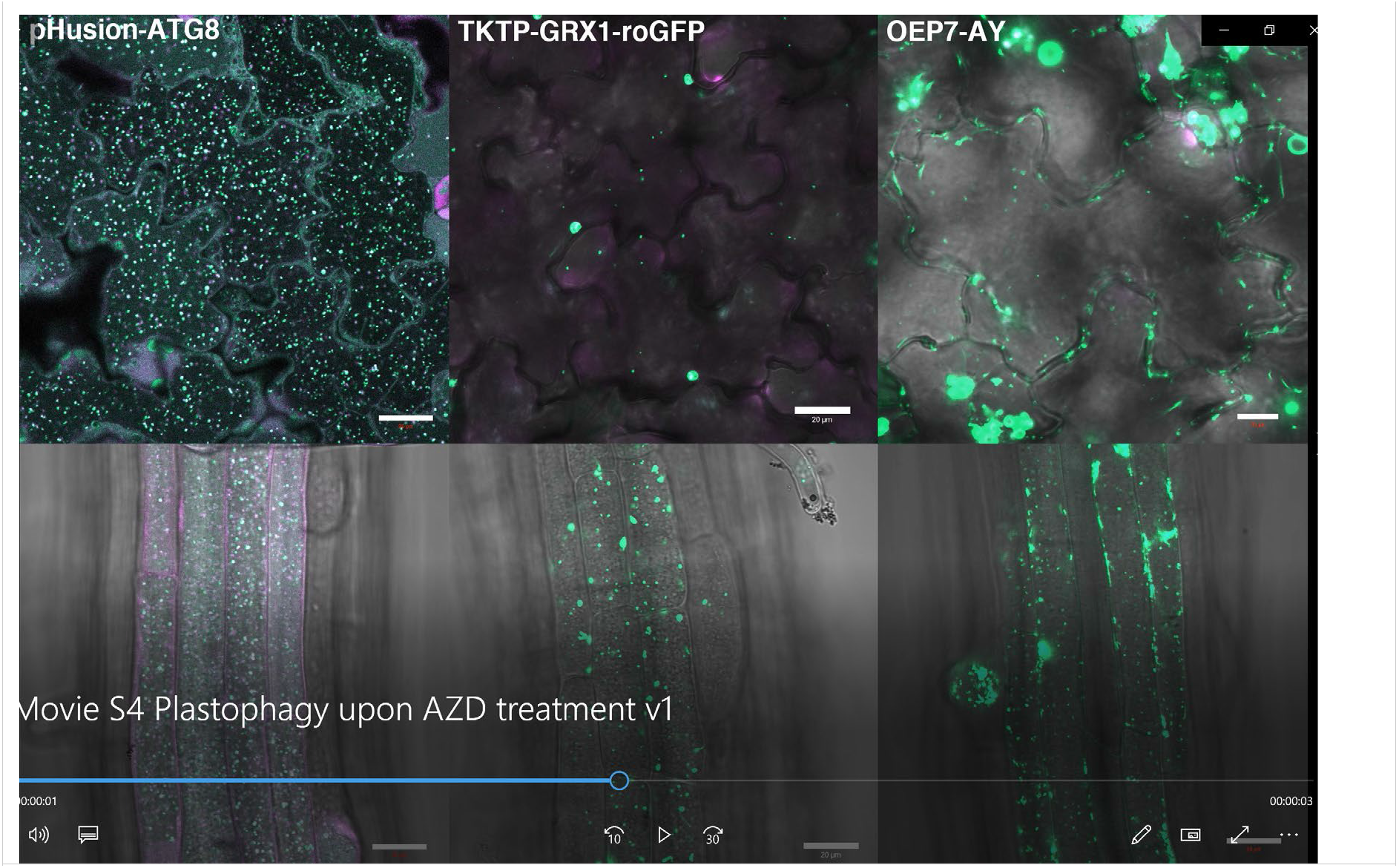
Plastid markers are detectable in a form of vacuolar puncta in the shoots and roots treated with AZD/ConA. Time-lapse confocal microscopy movies of shoot and root epidermal cells expressing fluorescent marker for plastid stroma (TKTP-GRX1-roGFP) or envelope marker (OE-YA). Seedlings were subjected to AZD/ConA treatment 24h prior to imaging. Seedlings expressing pHusion-ATG8 were used as a positive control for efficacy of autophagy induction. Scale bar, 20 μm.

**Movie S3.**
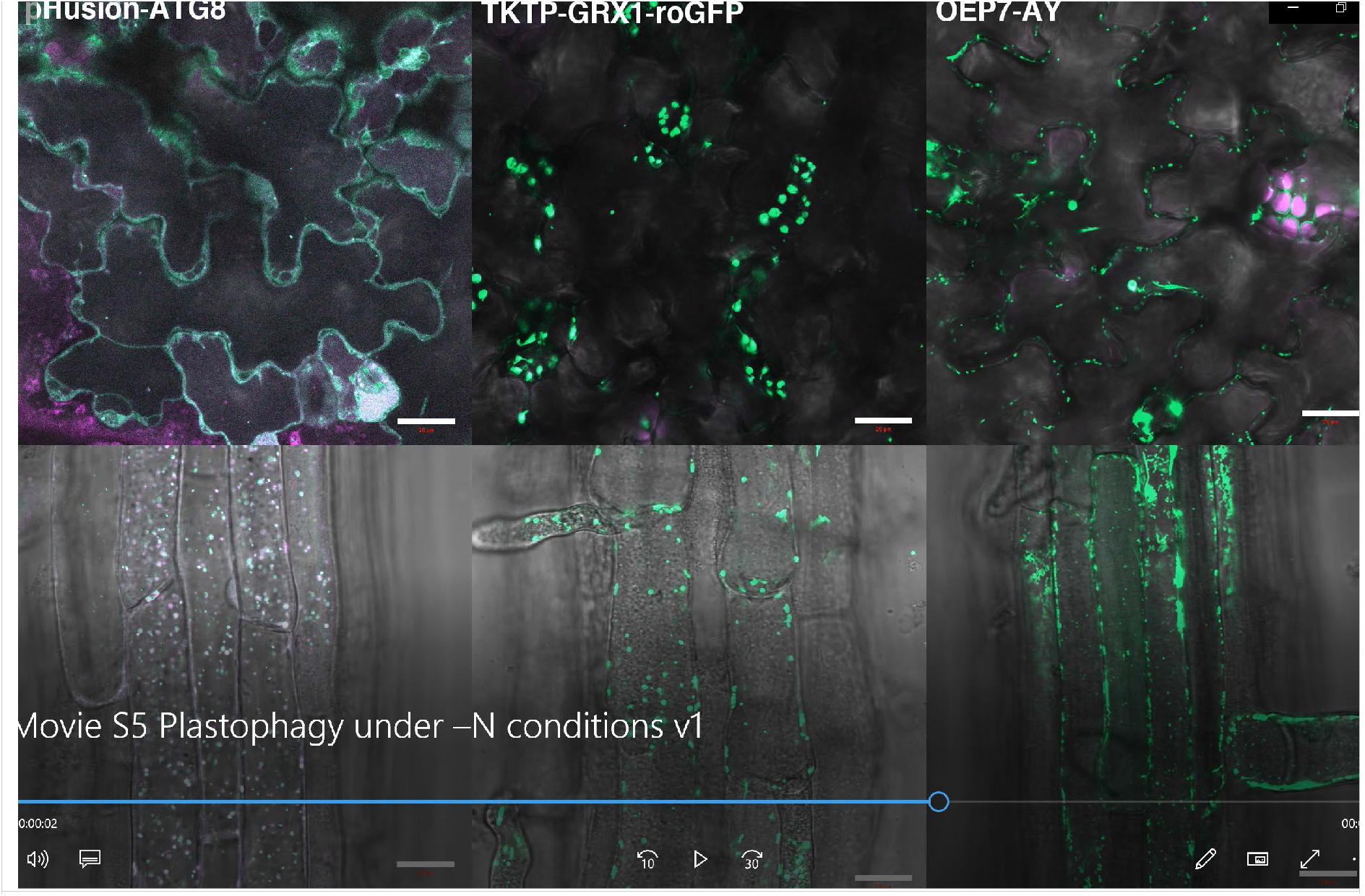
Both plastid markers were detectable in the vacuoles of roots under –N conditions, but only stromal marker was detectable in the vacuoles of shoots. Time-lapse confocal microscopy movies of shoot and root epidermal cells expressing fluorescent marker for plastid stroma (TKTP-GRX1-roGFP) or envelope marker (OE-YA). Seedlings were subjected to nitrogen-depleted conditions for 24h prior to imaging. Seedlings expressing pHusion-ATG8 were used as a positive control for efficacy of autophagy induction. Scale bar, 20 μm.

**Movie S4.**
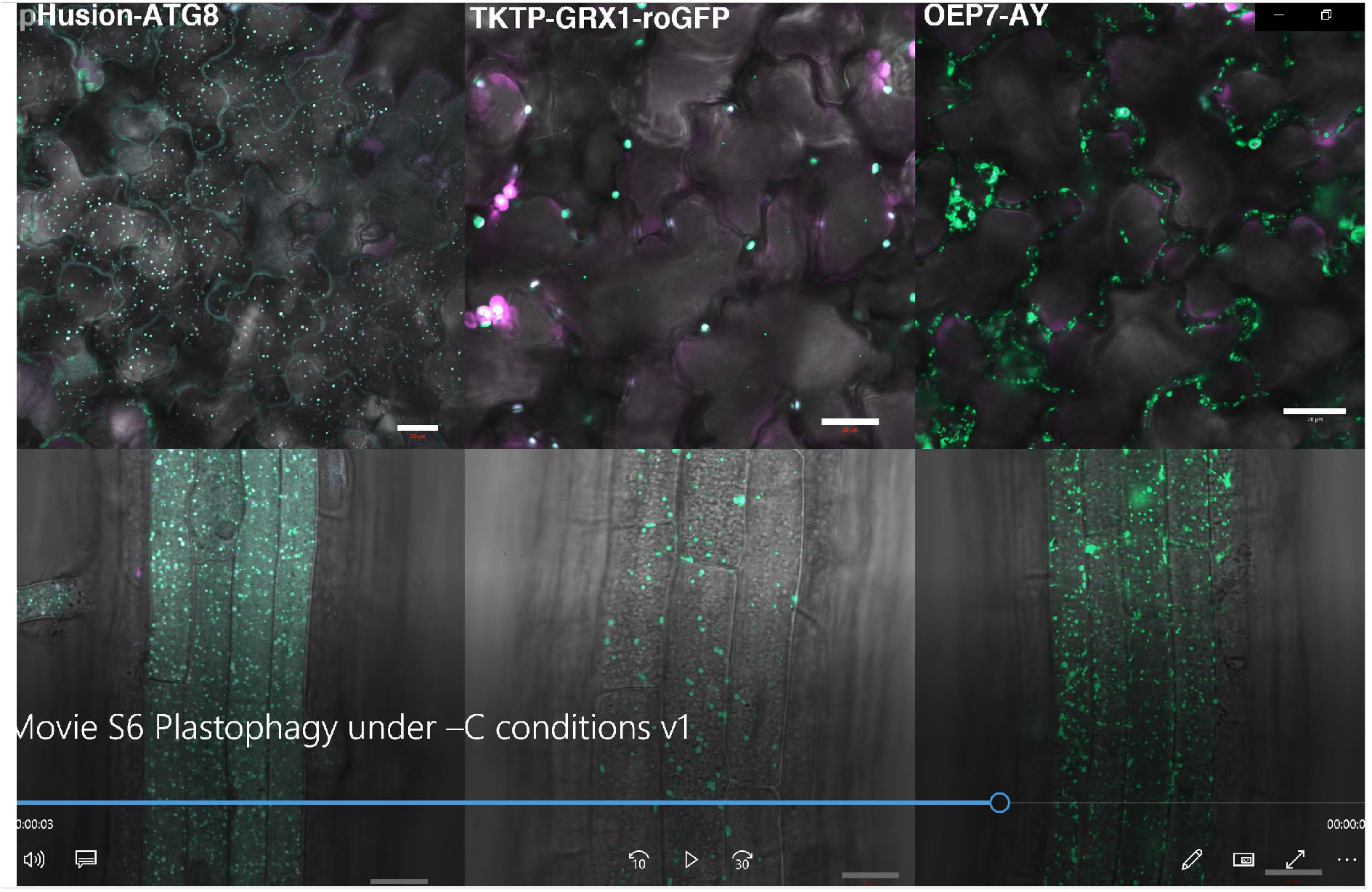
Both plastid markers were detectable in the vacuoles of roots under –C conditions, but only stromal marker was detectable in the vacuoles of shoots. Time-lapse confocal microscopy movies of shoot and root epidermal cells expressing fluorescent marker for plastid stroma (TKTP-GRX1-roGFP) or envelope marker (OE-YA). Seedlings were subjected to carbon-depleted conditions for 24h prior to imaging. Seedlings expressing pHusion-ATG8 were used as a positive control for efficacy of autophagy induction. Scale bar, 20 μm.

**Movie S5.**
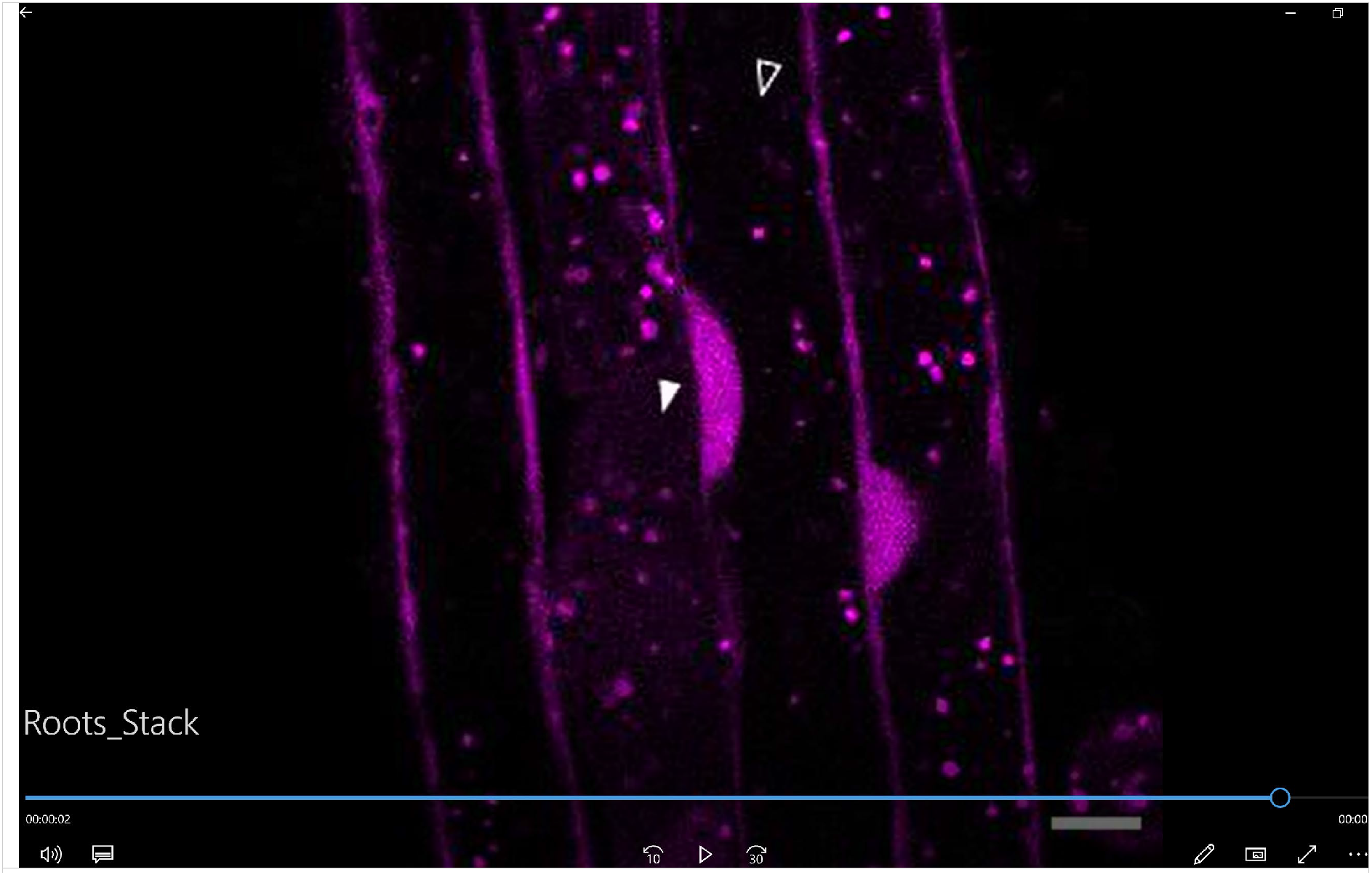
Illustrative examples of the differences between the appearances of autophagic bodies and cytoplasmic strands seen on an optical section of Arabidopsis root cells. Time-lapse confocal microscopy data obtained on the shoots of seedlings expressing pHusion-ATG8 and subjected to 24h of AZD/ConA treatment. Solid arrowheads indicate cytoplasmic strand and empty arrowheads point at autophagic bodies. Scale bar, 25 µ M

## References

1. Rosado-Souza, L., Yokoyama, R., Sonnewald, U. & Fernie, A. R. Understanding source–sink interactions: Progress in model plants and translational research to crops. Molecular Plant vol. 16 Preprint at 10.1016/j.molp.2022.11.015 (2023).

2. Jia, Z., Giehl, R. F. H. & Von Wirén, N. The Root Foraging Response under Low Nitrogen Depends on DWARF1-Mediated Brassinosteroid Biosynthesis 1[OPEN]. doi:10.1104/pp.20.00440.

3. Mizushima, N. A brief history of autophagy from cell biology to physiology and disease. Nat Cell Biol 20, 521–527 (2018).

4. Marshall, R. S. & Vierstra, R. D. Autophagy: The Master of Bulk and Selective Recycling. Annu Rev Plant Biol 69, 173–208 (2018).

5. Burkart, G. M. & Brandizzi, F. A Tour of TOR Complex Signaling in Plants. Trends in Biochemical Sciences vol. 46 Preprint at 10.1016/j.tibs.2020.11.004 (2021).

6. Pacheco, J. M. et al. The tip of the iceberg: Emerging roles of TORC1, and its regulatory functions in plant cells. Journal of Experimental Botany Preprint at 10.1093/jxb/eraa603 (2021).

7. Liu, G. Y. & Sabatini, D. M. mTOR at the nexus of nutrition, growth, ageing and disease. Nature Reviews Molecular Cell Biology vol. 21 Preprint at 10.1038/s41580-019-0199-y (2020).

8. Liu, Y. et al. Diverse nitrogen signals activate convergent ROP2-TOR signaling in Arabidopsis. Dev Cell 56, (2021).

9. Tsukada, M. & Ohsumi, Y. Isolation and characterization of autophagy-defective mutants of Saccharomyces cerevisiae. FEBS Lett 333, 169–174 (1993).

10. Yoshimoto, K. & Ohsumi, Y. Unveiling the Molecular Mechanisms of Plant Autophagy — From Autophagosomes to Vacuoles in Plants Special Focus Issue – Mini Review. Plant Cell Physiol 59, 1337–1344 (2018).

11. Adriaenssens, E., Ferrari, L. & Martens, S. Orchestration of selective autophagy by cargo receptors. Current Biology 32, R1357–R1371 (2022).

12. Zaffagnini, G. & Martens, S. Mechanisms of Selective Autophagy. J Mol Biol 428, 1714–1724 (2016).

13. Bieber, A. et al. In situ structural analysis reveals membrane shape transitions during autophagosome formation. (2022) doi:10.1073/pnas.

14. Onodera, J. & Ohsumi, Y. Ald6p Is a Preferred Target for Autophagy in Yeast, Saccharomyces cerevisiae. Journal of Biological Chemistry 279, 16071–16076 (2004).

15. Chresta, C. M. et al. AZD8055 is a potent, selective, and orally bioavailable ATP-competitive mammalian target of rapamycin kinase inhibitor with in vitro and in vivo antitumor activity. Cancer Res 70, (2010).

16. Montané, M. H. & Menand, B. ATP-competitive mTOR kinase inhibitors delay plant growth by triggering early differentiation of meristematic cells but no developmental patterning change. J Exp Bot 64, (2013).

17. Dauphinee, A. N. et al. Chemical screening pipeline for identification of specific plant autophagy modulators. Plant Physiol 181, 855–866 (2019).

18. Dauphinee, A., Ohlsson, J. & Minina, E. Tandem Tag Assay Optimized for Semi-automated in vivo Autophagic Activity Measurement in Arabidopsis thaliana roots. Bio Protoc 10, 1–16 (2020).

19. Guichard, M., Holla, S., Wernerová, D., Grossmann, G. & Minina, E. A. RoPod , a customizable toolkit for non-invasive root imaging , reveals cell type-specific dynamics of plant autophagy. bioRxiv 1–21 (2023).

20. Nazio, F. & Cecconi, F. Autophagy up and down by outsmarting the incredible ULK. Autophagy 13, 967–968 (2017).

21. Holczer, M. et al. Fine-tuning of AMPK-ULK1-mTORC1 regulatory triangle is crucial for autophagy oscillation. (2020) doi:10.1038/s41598-020-75030-8.

22. Klionsky, D. J., et al. Guidelines for the use and interpretation of assays for monitoring autophagy (3rd edition). Autophagy vol. 12 1554–8635 (Online) Preprint at 10.1080/15548627.2015.1100356 (2016).

23. Conway, L. J. & Poethig, R. S. Mutations of Arabidopsis thaliana that transform leaves into cotyledons. Proceedings of the National Academy of Sciences 94, 10209–10214 (1997).

24. Kellner, R. et al. ATG8 Expansion : A Driver of Selective Autophagy Diversi fi cation ? Trends Plant Sci 22, 204–214 (2017).

25. Holla, S. et al. ATG8 delipidation is dispensable for plant autophagy. Manuscript in preparation (2023).

26. Josse, E.-M. & Halliday, K. J. Skotomorphogenesis: The Dark Side of Light Signalling. Current Biology 18, R1144–R1146 (2008).

27. Bailey, B. N. et al. Modeling of Root Nitrate Responses Suggests Preferential Foraging Arises From the Integration of Demand, Supply and Local Presence Signals. (2020) doi:10.3389/fpls.2020.00708.

28. Campbell, R. E. et al. A monomeric red fluorescent protein. Proc Natl Acad Sci U S A 99, (2002).

29. Svenning, S., Lamark, T., Krause, K. & Johansen, T. Plant NBR1 is a selective autophagy substrate and a functional hybrid of the mammalian autophagic adapters NBR1 and p62/SQSTM1. Autophagy 7, 993–1010 (2011).

30. Jauh, G. Y., Phillips, T. E. & Rogers, J. C. Tonoplast intrinsic protein isoforms as markers for vacuolar functions. Plant Cell 11, (1999).

31. Krebs, M. et al. Arabidopsis V-ATPase activity at the tonoplast is required for efficient nutrient storage but not for sodium accumulation. Proc Natl Acad Sci U S A 107, (2010).

32. Ugalde, J. M. et al. Chloroplast-derived photo-oxidative stress causes changes in H2O2 and EGSH in other subcellular compartments. Plant Physiol 186, 125–141 (2021).

33. Mehlmer, N. et al. A toolset of aequorin expression vectors for in planta studies of subcellular calcium concentrations in Arabidopsis thaliana. J Exp Bot 63, 1751–1761 (2012).

34. Liu, F. et al. AUTOPHAGY-RELATED14 and its associated phosphatidylinositol 3-kinase complex promote autophagy in arabidopsis. Plant Cell 32, (2020).

35. Glab, N. et al. The impact of Arabidopsis thaliana SNF1-related-kinase 1 (SnRK1)-activating kinase 1 (SnAK1) and SnAK2 on SnRK1 phosphorylation status: characterization of a SnAK double mutant. (2016) doi:10.1111/tpj.13445.

36. Dong, S., Zhang, F. & Beckles, D. M. A Cytosolic Protein Kinase STY46 in Arabidopsis thaliana Is Involved in Plant Growth and Abiotic Stress Response. doi:10.3390/plants9010057.

37. Nakatogawa, H. Two ubiquitin-like conjugation systems that mediate membrane formation during autophagy. Autophagy: Molecules and Mechanisms 55, 39–50 (2013).

38. Schreiber, A. et al. Multilayered regulation of autophagy by the Atg1 kinase orchestrates spatial and temporal control of autophagosome formation. Mol Cell 81, (2021).

39. Dobrenel, T. et al. TOR Signaling and Nutrient Sensing. Annu Rev Plant Biol 67, 261–285 (2016).

40. Zhou, Z. et al. Phosphorylation regulates the binding of autophagy receptors to FIP200 Claw domain for selective autophagy initiation. Nat Commun 12, (2021).

41. Ran, J., Hashimi, S. M. & Liu, J. Z. Emerging roles of the selective autophagy in plant immunity and stress tolerance. International Journal of Molecular Sciences vol. 21 Preprint at 10.3390/ijms21176321 (2020).

42. Zess, E. K. et al. N-terminal β-strand underpins biochemical specialization of an ATG8 isoform. PLoS Biol 17, e3000373 (2019).

43. Mhimdi, M. & Pérez-Pérez, J. M. Understanding of Adventitious Root Formation: What Can We Learn From Comparative Genetics? Frontiers in Plant Science vol. 11 Preprint at 10.3389/fpls.2020.582020 (2020).

44. Wu, F.-H. et al. Tape-Arabidopsis Sandwich-a simpler Arabidopsis protoplast isolation method. Plant Methods 5, 1–10 (2009).

45. Robert, S., Zouhar, J., Carter, C. & Raikhel, N. Isolation of intact vacuoles from Arabidopsis rosette leaf-derived protoplasts. Nat Protoc 2, (2007).

